# Loss of Nuclear TDP-43 Impairs Lipid Metabolism in Microglia-Like Cells

**DOI:** 10.1101/2025.09.17.676815

**Authors:** Khushbu Kabra, Dallin Dressman, Ryan Talcoff, Maedot Yidenk, Olivia M. Rifai, Benjamin N. Hoover, Neil A. Shneider, Wassim Elyaman, Estela Area-Gomez, Elizabeth M. Bradshaw

**Author notes:** Correspondence should be addressed to E.M.B.

## Abstract

Amyotrophic lateral sclerosis (ALS) is a fatal neurodegenerative disease marked by progressive motor neuron loss, with TDP-43 pathology present in over 90% of cases. While neuroinflammation is a recognized hallmark, the role of microglia in ALS pathogenesis remains incompletely understood. Here, we demonstrate that TDP-43 regulates microglial function via triglyceride metabolism. Using shRNA-mediated *TARDBP* knockdown in human monocyte-derived microglia-like cells (MDMi), we observed suppressed cholesterol biosynthesis, upregulation of fatty acid metabolism genes, lipid droplet accumulation, enhanced phagocytic activity, and increased IL-1β production. Inhibiting diacylglycerol acyltransferase (DGAT) enzymes reduced lipid droplet formation, phagocytosis, and IL-1β, directly linking the triglyceride pathway to microglial activation. Patient-derived MDMi from both sporadic and *TARDBP*-mutant ALS cases showed overlapping as well as distinct alterations, some of which were reversed by DGAT inhibition. Our findings identify dysregulated triglyceride metabolism as a novel pathway through which TDP-43 mediates microglial dysfunction, highlighting a potential therapeutic target for ALS.

**Graphical Abstract:** 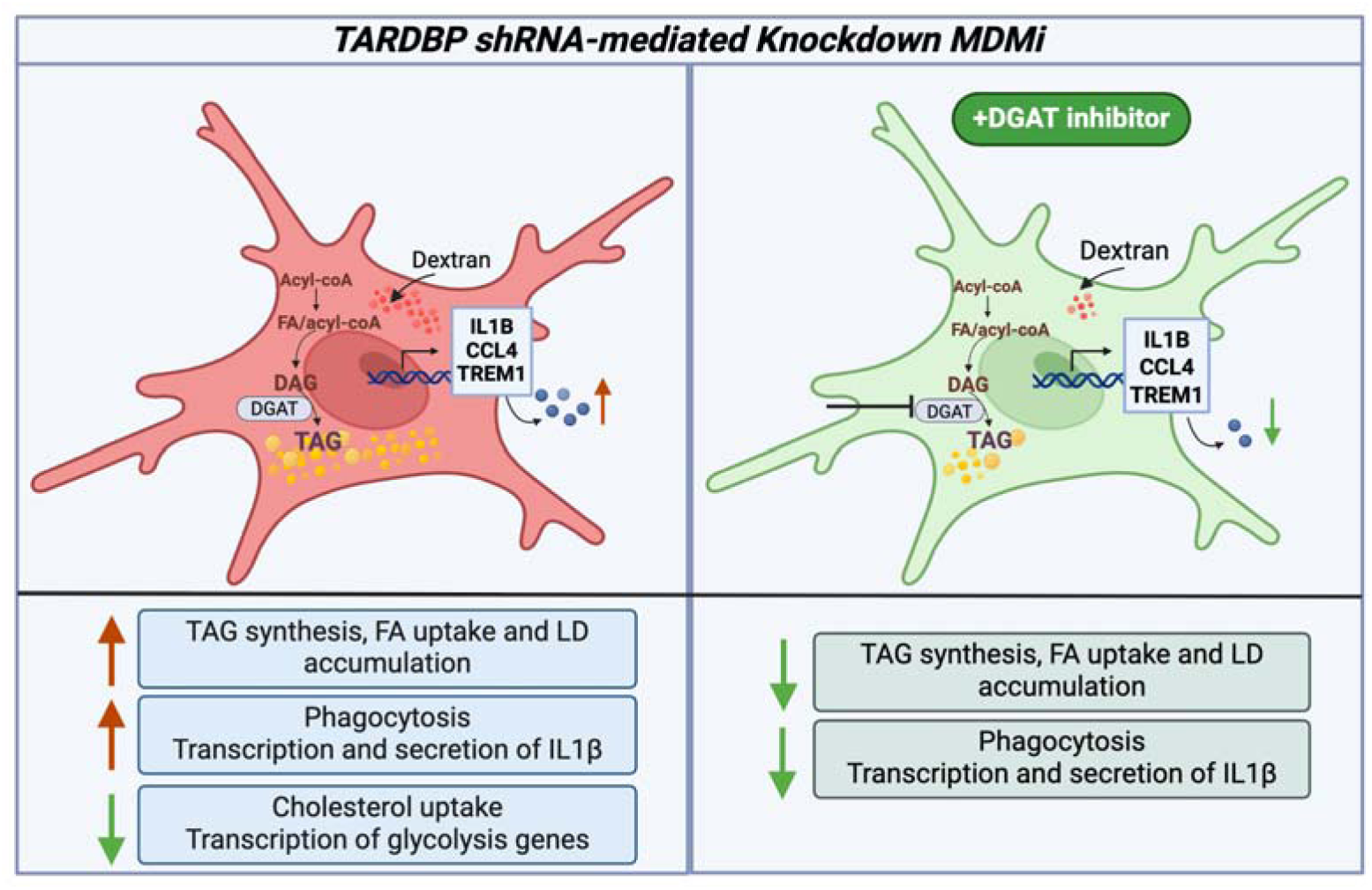

**Highlights:**

- TDP-43 nuclear depletion causes increased LD, driven by triglyceride accumulation.
- TDP-43 nuclear depletion causes increased phagocytosis and pro-inflammatory cytokine expression.
- Inhibiting triglyceride synthesis using DGAT inhibitors rescues LD and pro-inflammatory phenotype in TDP-43 depleted MDMi
- ALS patient-derived MDMi display increased LD and *IL1B* expression, rescued by DGAT inhibitors

## Introduction

Amyotrophic lateral sclerosis (ALS) is a progressive neurodegenerative disorder primarily affecting motor neurons. Although only about 10% of cases are familial, an increasing number of genes with diverse biological functions have been linked to both familial and sporadic forms of ALS [1,2]. Multiple cellular pathways, including protein degradation, mitochondrial dysfunction, and autophagy, to name a few, have been implicated in disease pathogenesis, yet the precise mechanisms remain poorly understood. Despite this genetic and mechanistic heterogeneity, common features such as metabolic dysfunction, neuroinflammation, and microglial dysregulation consistently emerge across ALS cases [3–5].

Microglia are key innate immune cells in the central nervous system (CNS), known to be dysregulated in various neurodegenerative diseases, including ALS [6–8]. Although the presence of neuroinflammation, including activation of glial cells and infiltration of peripherally-derived innate and adaptive immune cells, is a consistent hallmark of ALS [9], the specific mechanism by which microglia contribute to disease pathology is unclear [10]. Microglia sequencing studies in ALS have demonstrated significant, distinct alterations in gene expression compared to both healthy subjects and individuals with other neurodegenerative diseases [11–13]. Transcriptomic studies of post-mortem spinal cord tissues implicate glial activation in ALS [14,15]. In particular, chronic or excessive microglial activation has been proposed as a mechanism driving the transition from neuroprotective to neurotoxic phenotypes, making microglia a potential therapeutic target in ALS [16,17].

An important characteristic of microglia is that they are highly mobile and dynamic, constantly modifying their shape and membrane structure to survey the environment and respond to pathogens and injury [18]. Lipids have a crucial role in regulating membrane structure and are therefore important in microglia function [19]. Additionally, lipid microdomains are required for the localization of signaling receptors and regulating immune pathways [20–22]. Microglia are also known to be metabolically altered in many neurodegenerative diseases.

Notably, lipid droplets (LDs), which store cholesterol esters, triglycerides, and other lipids within cells [23], have been shown to accumulate in aging as well as Alzheimer’s disease (AD) [24]. Interestingly, the ApoE4 genotype, which confers significant risk for AD, has been linked to lipid droplet formation in induced pluripotent stem cell (iPSC)-derived microglia. These ‘lipid-laden’ microglia were enriched in cellular senescence genes and shown to drive neurotoxicity [25]. It is also known that lipopolysaccharide (LPS) stimulation in microglia causes accumulation of LDs, which activate inflammatory pathways in the brain that can become neurotoxic over time and contribute to neuroinflammation [26]. The mechanisms underlying microglial LD formation in different contexts remain elusive. Studies have shown metabolic dysregulation, specifically lipid alterations, to be a consistent feature in ALS, with numerous studies demonstrating alterations in lipid pathways in both patient biofluids and disease models [27–29]. However, the mechanisms regulating lipid dysfunction in ALS, specifically in microglia, have not been well-studied [5,30].

*TARDBP*, which encodes the TAR DNA-binding protein 43 (TDP-43), is a key gene implicated in ALS pathology [31]. TDP-43 aggregation and nuclear depletion are hallmarks in ∼95% of ALS cases [32]. Previous studies in oligodendrocytes, motor neurons, as well as HeLa and HEK293 cell lines have demonstrated the role of TDP-43 in regulating lipid metabolism, including cholesterol biosynthesis and efflux pathways, but the mechanisms by which these disruptions impact ALS remain unclear. A recent study showed that overexpression of *TARDBP* in HEK293 cells resulted in defects in cholesterol biosynthesis [33], whereas another study showed that conditional knockout of *TARDBP* in mouse oligodendrocytes suppressed cholesterol biosynthesis gene expression [34]. Although these studies implicate a role for TDP-43 in regulating cholesterol biosynthesis via SREBP2, the underlying mechanisms remain to be established, as does the impact on immune cells like microglia, and the contribution to ALS pathology.

Here, we investigated the impact of *TARDBP* knockdown on lipid metabolism and microglial function using human MDMi. Previous work has demonstrated that MDMi generated from ALS patient-derived blood display nuclear loss of TDP-43 and exhibit other ALS phenotypes such as altered cytokine expression and function [35]. We therefore used a *TARDBP* shRNA-mediated knockdown model to specifically investigate lipid alterations and determine how loss of TDP-43 impacts microglia function. Additionally, we analyzed patient-derived MDMi from both sporadic ALS (sALS) cases and patients with *TARDBP* mutations (TDP-ALS) and observed both shared and distinct features when compared to the shRNA knockdown model.

## Results

### TARDBP knockdown causes nuclear TDP-43 depletion in MDMi

To investigate how TDP-43 depletion alters lipids in a microglia model, we employed shRNA-mediated gene knockdown in MDMi, achieving an average *TARDBP* knockdown efficiency of approximately 70% (**Fig. 1A**). The demographic information of donors used can be found in **Supplementary Table 1**. Knockdown efficiency was validated by both immunocytochemistry (**Fig. 1B**) and Western blot analysis (**Fig. S1A, B**). Confocal imaging showed a 40% reduction in mean nuclear TDP-43 intensity (**Fig. 1C**) and a significant reduction in the nuclear to cytoplasmic ratio of TDP-43 staining (**Fig. S1C**), but no significant change in cytoplasmic TDP-43 (**Fig. 1D**). We also stained for phospho-TDP-43 (pTDP-43), given that pTDP-43 is known to increase in ALS and is a marker for increased cytosolic TDP-43 accumulation [36,37]. However, there were no significant alterations in nuclear or cytoplasmic pTDP-43 (**Fig. S2 A-C**), or the nuclear to cytoplasmic ratio of pTDP-43 (**Fig. S2D**).

**Fig 1:**
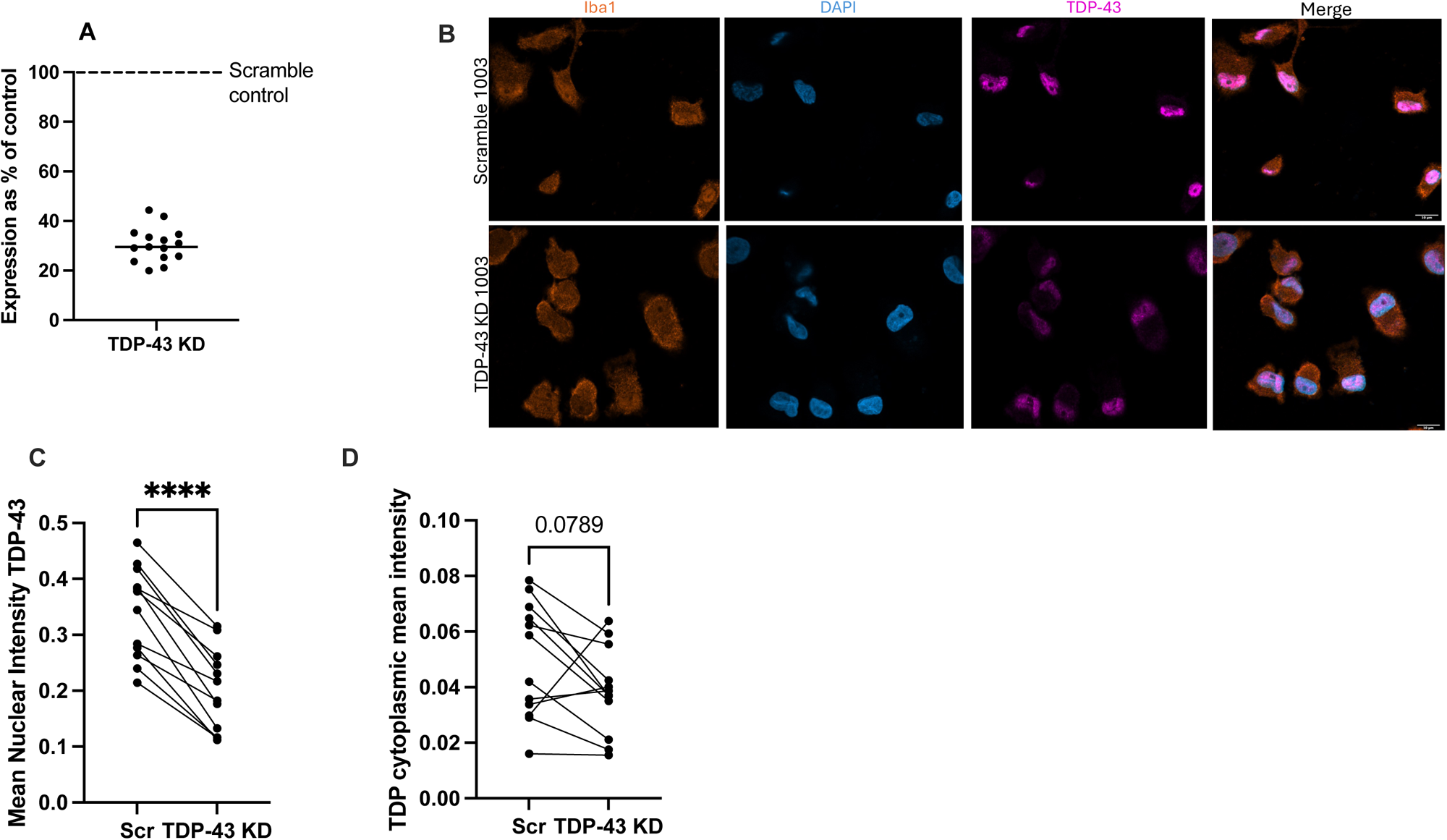
*TARDBP* knockdown results in TDP-43 nuclear depletion in MDMi. **A**) Expression of *TARDBP* in knockdown (TDP-43 KD) samples as a percentage of the scramble control (Scr) (N=15) measured using qPCR. **B**) 63X Confocal imaging showing staining for Iba1, DAPI and TDP-43; the last panel shows a merge. Scale bar = 10μm. **C**) Quantification of nuclear TDP-43 stain from CellProfiler images, performed using CellProfiler (N=12). **D**) Quantification of cytoplasmic TDP-43 stain from CellProfiler images, performed using CellProfiler (N=12). Each connected line shows paired samples from the same donor. Statistical analysis: Paired t-test (**** = p<0.0001)

### TARDBP knockdown significantly alters lipid metabolism and immune genes in MDMi

To assess the impact of *TARDBP* knockdown (TDP-43 KD), we used the Fluidigm Biomark microfluidics system to analyze gene expression with a targeted panel of 110 genes related to lipid metabolism (including cholesterol, phospholipid/sphingolipid, triglyceride, and fatty acid metabolism), glycolysis, and immune function, and compared results to scramble controls. STRING analysis [38] shows the genes in three main clusters as expected, immune function, lipid metabolism, and glycolysis, although many of the genes have related functions (**Fig. 2A**). As shown in the volcano plot (**Fig. 2B**), many genes were significantly altered in the knockdown. Interestingly, of the 110 genes measured, only 23 were upregulated. 15 of these reached statistical significance with an unadjusted p-value<0.05, while only 2 were significant with an adjusted p-value<0.05 (**Fig. 2C**). Many of these genes were related to immune function, with pro-inflammatory cytokines *CCL4* and *IL1*β being most highly upregulated (**Fig. 2C**). *TREM1*, associated with increased inflammatory phenotypes, was also highly upregulated. Among the lipid cluster, fatty acid-associated genes like *FABP4*, *ELOVL3,* and *BSCL2* were upregulated. These genes are involved in fatty acid uptake and transport, elongation, and lipid droplet formation [39–42].

**Fig 2:**
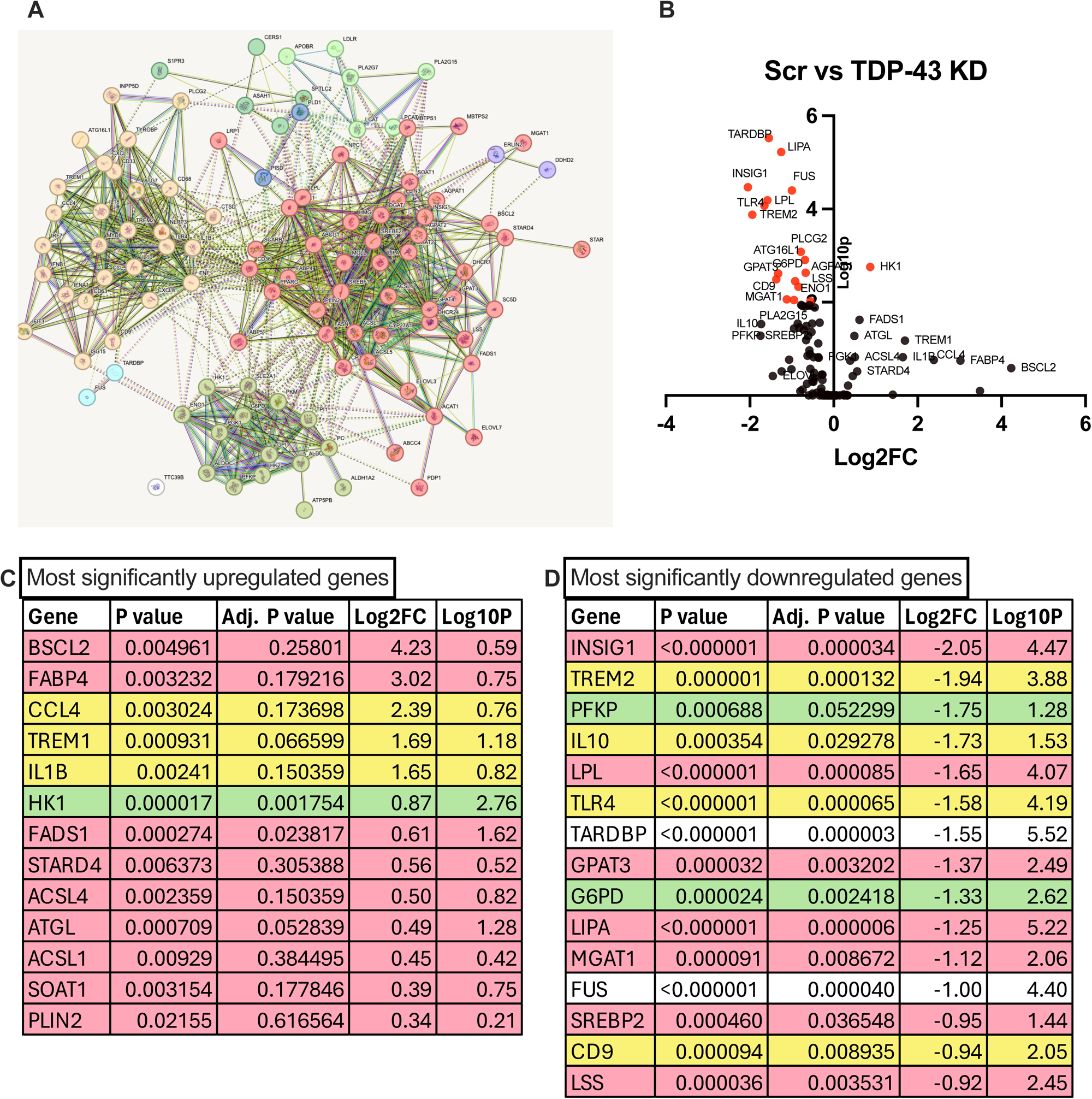
*TARDBP* knockdown significantly alters lipid metabolism gene expression. **A**) STRING analysis of Fluidigm genes showing 3 clusters of genes included in the panel: lipid metabolism (red), glycolysis (green) and immune function (yellow). **B**) Volcano plot for Fluidigm comparing gene expression between TARDBP knockdown (TDP-43 KD) and scramble control (Scr)(N=15). **C**) Table showing most significantly upregulated genes with unadjusted p-values <0.05 **D**) Table showing 15 most significantly downregulated genes with adjusted p-values <0.05 Statistical analysis: Multiple paired t-test corrected for multiple hypothesis testing using Holm-Sidak method. Adjusted p-values used for volcano plot.

Of the 110 analyzed genes, 87 were downregulated, with 59 showing statistically significant changes with an unadjusted p-value<0.05, while 33 of those were significant with an adjusted p-value<0.05. Among the top 15 downregulated genes, *INSIG1,* which encodes the protein INSIG1 that binds to and regulates SREBP2, exhibited the greatest reduction in expression (Log2FC= -2) (**Fig. 2D**). This is consistent with a previous study in mouse oligodendrocytes where conditional knockout of *TARDBP* resulted in a significant reduction in *INSIG1* [34]. Other genes related to *INSIG1,* such as *SREBP2* and *MBTPS1,* were also downregulated in our study, suggesting suppression of cholesterol biosynthesis pathways, again consistent with previous studies [34,43]. Other significantly downregulated genes included *TREM2* and *LPL,* both highly expressed in microglia, and known to be involved in neurodegenerative diseases like AD through their function in regulating microglia lipid homeostasis as well as immune function [44,45]. *TREM2* expression has been found to be increased in spinal cords of SOD1-ALS mice and in reactive microglia from ALS postmortem tissues [13,46]. The Alzheimer’s disease-associated genetic variation TREM2 R47H, which modulates ligand binding, has also been implicated in ALS, supporting the importance of correctly functioning lipid metabolism in microglia [47,48].

We confirmed the increase in *IL1*β expression by qPCR and ELISA and found that it was indeed significantly upregulated at both the gene and protein level (**Fig. S3A, B**). We also measured soluble TREM2 (a biologically active fragment of TREM2) and found a significant reduction in the *TARDBP* knockdown (**Fig. S3C**). In addition to *TREM2*, *IL10* gene expression was significantly decreased, suggesting a possible downregulation of anti-inflammatory pathways while pro-inflammatory pathways are upregulated (**Fig. 2D**).

Interestingly, genes related to glycolysis (*PFKP* and *G6PD)* were among those most significantly downregulated in the knockdown (**Fig. 2D**), suggesting alterations in glucose metabolism that could drive a metabolic shift, which is a known feature of ALS [49,50].

### TDP-43 depletion causes increased lipid droplet accumulation in MDMi

LD accumulation has emerged as a significant phenotype in diseased/abnormal microglia and has been associated with AD, tauopathies, and aging [24,25,51], although the pathomechanism is not clear. Given the role of LD accumulation as a marker for microglia dysfunction, as well as significant alterations of genes like *TREM2, LPL, BSCL2,* and *FABP4* in our knockdown, we sought to determine whether TDP-43-depleted MDMi have altered LD accumulation. Using LipidTox green to stain for neutral lipids (usually stored in lipid droplets), we found a significant increase in LipidTox intensity in the *TARDBP* knockdown cells compared to controls (**Fig. 3A, B**), pointing to altered lipid metabolism and bioenergetics.

**Fig 3:**
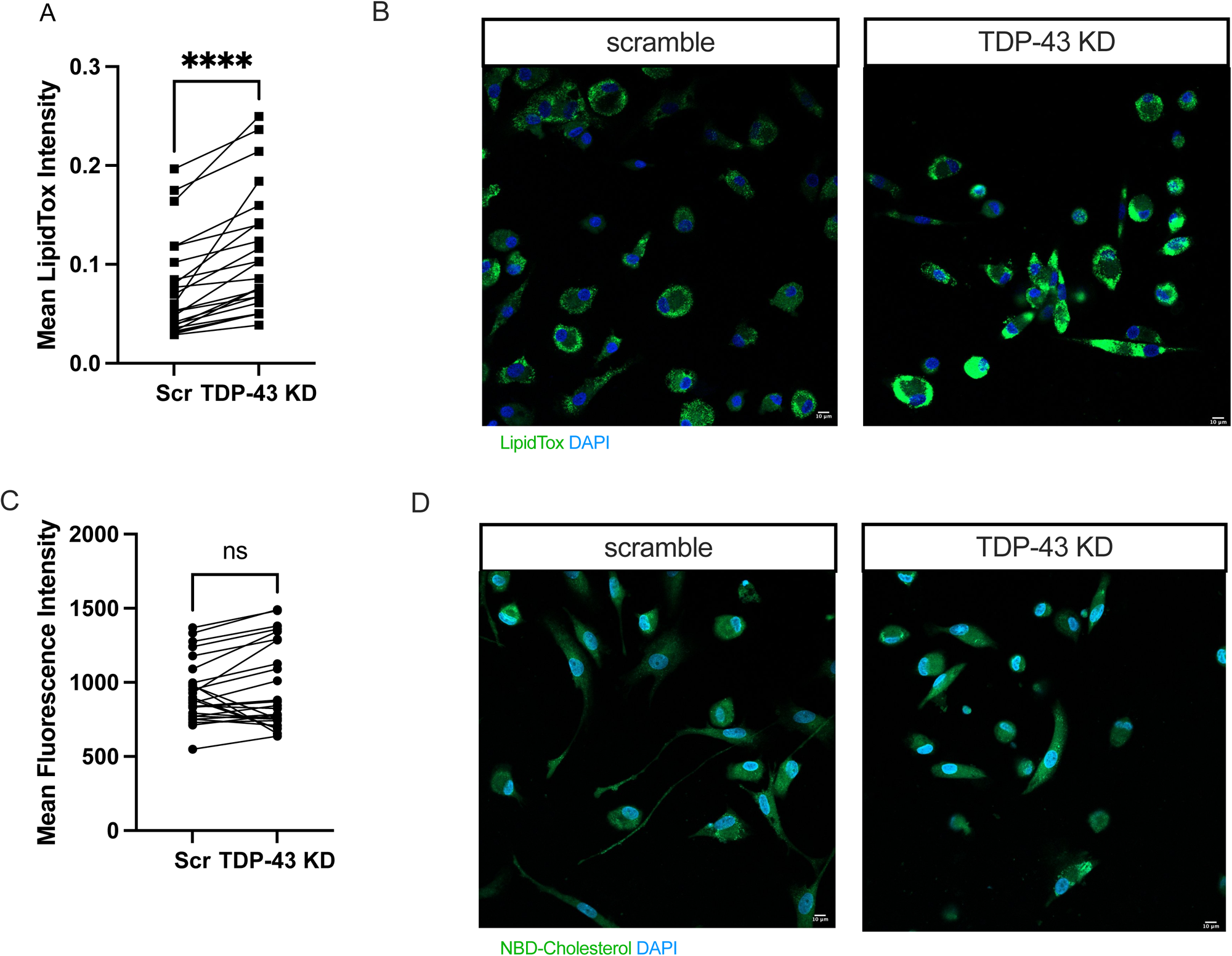
TDP-43 depletion causes increased lipid droplet accumulation in MDMi. **A**) Quantification of confocal imaging of LipidTox staining using CellProfiler (N= 23) for scramble control (Scr) and *TARDBP* knockdown (TDP-43 KD). **B**) Confocal imaging of LipidTox stain comparing scramble control (Scr) and *TARDBP* knockdown (TDP-43 KD), 20X. Scale bar = 10μm. **C)** NBD-cholesterol uptake fluorescence intensity quantified by plate reader (N=25). **D**) Confocal imaging of NBD-cholesterol comparing scramble control and TDP-43 KD, 20X. Scale bar = 10μm. Statistical analysis: Paired t-test.

### TDP-43 depletion does not alter cholesterol uptake in MDMi

Next, we investigated whether the accumulation of LDs was driven by changes in cholesterol uptake or efflux, given the dramatic alterations we observed in the expression of cholesterol-associated genes upon *TARDBP* knockdown.

Under conditions of low intracellular cholesterol levels, INSIG1 dissociates from the SCAP-SREBP2 complex, allowing SREBP2 to translocate to the endoplasmic reticulum (ER) from the Golgi and activate the transcription of various cholesterol biosynthesis genes, including HMGCR [52]. Downregulation of these genes, as observed in our *TARDBP* knockdown, may impair the cell’s capacity to activate cholesterol biosynthesis and could increase external uptake of cholesterol. Using a fluorescently labeled analogue of cholesterol (NBD-cholesterol), which has been previously shown to mimic cholesterol uptake via lipoproteins in cells [53–55], we measured the uptake of cholesterol in MDMi. Interestingly, we found no difference in NBD-cholesterol uptake between controls and *TARDBP* knockdown cells (**Fig. 3C, D**). To verify that this assay does in fact recapitulate how cells physiologically uptake cholesterol from the media, we repeated the experiment in the presence of LPS stimulation, as other studies have shown that LPS treatment induces cholesterol uptake in immune cells [56]. Indeed, we saw that with LPS stimulation, both scramble control and *TARDBP* knockdown cells exhibit an increase in NBD-cholesterol uptake; however, this increase was only significant in scramble controls (**Fig. S4A, B**).

When comparing the ratio of NBD-cholesterol uptake in LPS-stimulated to unstimulated conditions, we observed significantly lower uptake in the *TARDBP* knockdown condition compared to the scramble control, indicating a reduced response to LPS (**Fig. S4C**). These findings suggest that although *INSIG1 and SREBP2* expression are decreased and cholesterol synthesis may be depressed, TDP-43 depleted MDMi do not increase cholesterol uptake. Additionally, when stimulated with LPS, TDP-43 depleted MDMi do not have a significant induction of cholesterol uptake as seen in the control. Gene expression of *SCARB1*, *LDLR*, and *LRP1*, receptors involved in lipoprotein uptake, was also downregulated in the *TARDBP* knockdown, which could explain why these cells are unable to upregulate cholesterol uptake. Taken together, these data suggest that the lipid droplet accumulation observed in our *TARDBP* knockdown is not driven by cholesterol uptake.

### TARDBP knockdown in MDMi reduces cellular total and free cholesterol levels

Our gene expression analysis also revealed downregulation of *ABCA1* and *ABCC4*, suggesting possible impairment of cholesterol efflux. Indeed, previous reports have shown that *ABCA1* deletion causes an accumulation of cholesterol esters as the cell is unable to efflux excess cholesterol [57–59]. To determine whether the lipid droplet accumulation observed in our *TARDBP* knockdown cells is a result of cholesterol ester accumulation, we measured total cholesterol (TC), free cholesterol (FC), and cholesterol esters (CE) in cells and supernatants of scramble and *TARDBP* knockdown MDMi. Cells are known to uptake free cholesterol from lipoproteins, which are esterified by the enzyme Acyl-coenzyme A: cholesterol acyltransferase-1 (ACAT1) to cholesterol esters, for storage in LDs.

We found that *TARDBP* knockdown cells exhibited decreased TC and FC, with no significant change in CEs (**Fig. 4A-C**). The supernatants showed a similar trend, with TC and FC levels being significantly reduced while CE did not change (**Fig. 4D-F**). The overall reduction in TC and FC suggests reduced cholesterol synthesis in *TARDBP* knockdown cells, with a concomitant decrease in efflux. We found no significant difference in the ratio of FC to TC in scramble compared to *TARDBP* knockdown cells and supernatants, which supports the idea that cholesterol esterification is not driving lipid droplets in these cells (**Fig. S4D-E**), as increased esterification would reduce the proportion of FC. This is also supported by the NBD-cholesterol uptake data, where we found no significant changes in uptake between scramble and knockdown.

**Fig 4:**
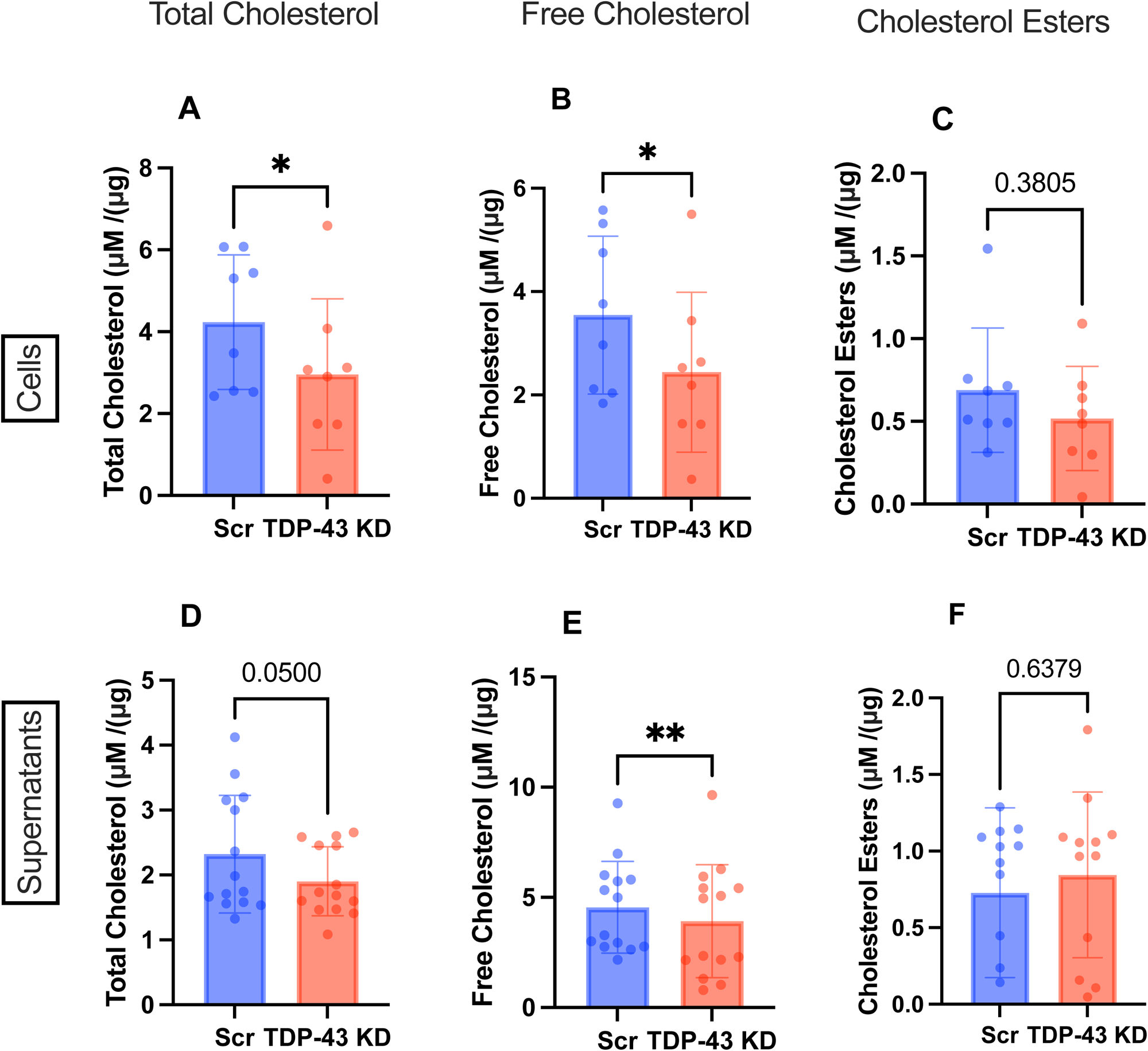
TDP-43 depletion does not alter cholesterol uptake in MDMi. **A-C**) Total cholesterol, Free Cholesterol and Cholesterol Ester levels in cells comparing scramble control (Scr) and *TARDBP* knockdown (TDP-43 KD) (N=8). **D-F**) Total cholesterol, Free Cholesterol and Cholesterol Ester levels in supernatants comparing scramble control (Scr) and *TARDBP* knockdown (TDP-43 KD) (N=14). All values were normalized using protein values calculated by Bradford analysis. Statistical analysis: paired t-test

To further confirm that the lipid droplet phenotype observed in *TARDBP* knockdown cells is not a result of cholesterol ester accumulation, we treated the scramble and knockdown MDMi with an ACAT1 inhibitor, which has previously been shown to reduce free cholesterol and cholesterol esters in cells [60]. As expected, ACAT1 inhibition led to a significant reduction in lipid droplets in scramble control cells; however, no such reduction was observed in TDP-43– depleted MDMi (**Fig. S4F**), again supporting the above finding that cholesterol esterification is not driving LD accumulation in these cells.

### TDP-43 depleted MDMi show increased glycerol and triglyceride levels

Lipid droplets are known to be comprised of both cholesterol esters and triglycerides [23,61]. We therefore measured triglyceride levels in these cells to interrogate whether triglyceride, rather than cholesterol ester accumulation, could be contributing to the lipid droplet phenotype observed in *TARDBP* knockdown cells. Triglyceride (TG) levels are calculated by subtracting the difference between total glycerol and free glycerol (which can be used to synthesize TGs). Interestingly, TGs were significantly increased in *TARDBP* knockdown cells, as were total glycerol levels, whereas free glycerol was not significantly altered (**Fig. 5A-C**).

**Fig 5:**
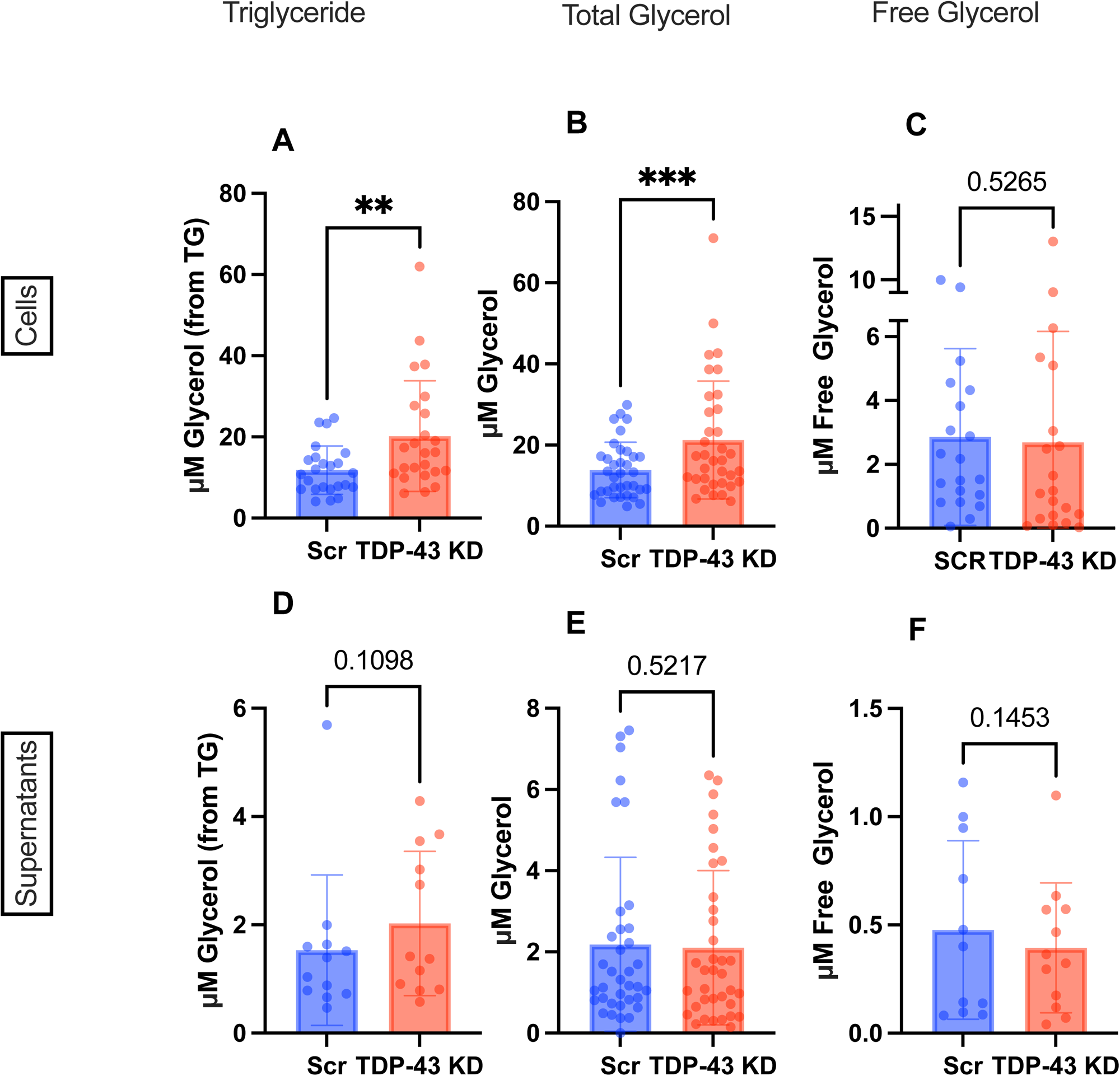
TDP-43 depletion causes triglyceride accumulation in MDMi. **A-C**) Triglycerides, total glycerol and free glycerol levels in scramble control (Scr) vs *TARDBP* knockdown (TDP-43 KD) in cell lysates measured using Promega TriGlo Assay. **D-F**) Triglycerides, total glycerol and free glycerol levels in scramble control (Scr) vs *TARDBP* knockdown (TDP-43 KD) in supernatants measured using Promega TriGlo Assay. (For cells: Triglyceride only n=24, for Total Glycerol N=35, for Free Glycerol (N=20). For sups: Triglyceride only N=12, Total Glycerol N=35, Free Glycerol N=12). Statistical analysis: paired t-test. Sample numbers differ for cells vs supernatants due to kit limitations.

No significant alterations were observed in the supernatants (**Fig. 5D-F**). However, the cell-to-supernatant ratio of total glycerol and TGs was significantly increased in the knockdown (**Fig. S5A-C**), while free glycerol remained unchanged. The increase in TGs and total glycerol, with stable free glycerol, suggests enhanced TG synthesis or possible defects in lipolysis.

### DGAT1/2 inhibitors reduce lipid droplets in TARDBP knockdown MDMi

Microglia have been shown to contain higher levels of triglycerides compared to other brain cell types, specifically astrocytes and neurons [62]. Additionally, a recent study found that triglyceride metabolism may be key in regulating microglia inflammatory pathways [63]. In this study, inhibitors of DGAT 1 and 2 (diacylglycerol acyltransferases 1 and 2), which control the rate-limiting step of triglyceride biosynthesis from diacylglycerol and fatty acids, were used to reduce triglyceride accumulation in iPSC-derived microglia. Given the possibility of increased triglyceride synthesis in our *TARDBP* knockdown model, we used a similar approach to determine whether DGAT inhibitors could reverse the lipid droplet phenotype observed.

First, we measured the effect of the DGAT inhibitors on glycerol and TG levels. Unsurprisingly, we found that DGAT inhibitors reduced total glycerol, free glycerol, and triglycerides in the *TARDBP* knockdown condition (**Fig. 6A-C**). However, no significant changes were seen in the scramble control, suggesting that TG accumulation, and possibly an increase in the activity of DGAT 1 and 2 enzymes, only occurs when nuclear TDP-43 levels are reduced. Interestingly, total glycerol levels were increased in the supernatants of DGAT inhibitor-treated samples, both in scramble and *TARDBP* knockdown conditions (**Fig. 6D**), suggesting that inhibiting triglyceride synthesis either reduces the incorporation of glycerol from the media, or increases the secretion of excess glycerol. As an additional control, we measured glycerol and TG levels in cells treated with ACAT1 inhibitors. As expected, ACAT1 inhibition did not alter total glycerol or TG levels in the scramble or knockdown cells, and there was no change in total glycerol levels in the supernatants (**Fig. S6A-D**).

**Fig 6:**
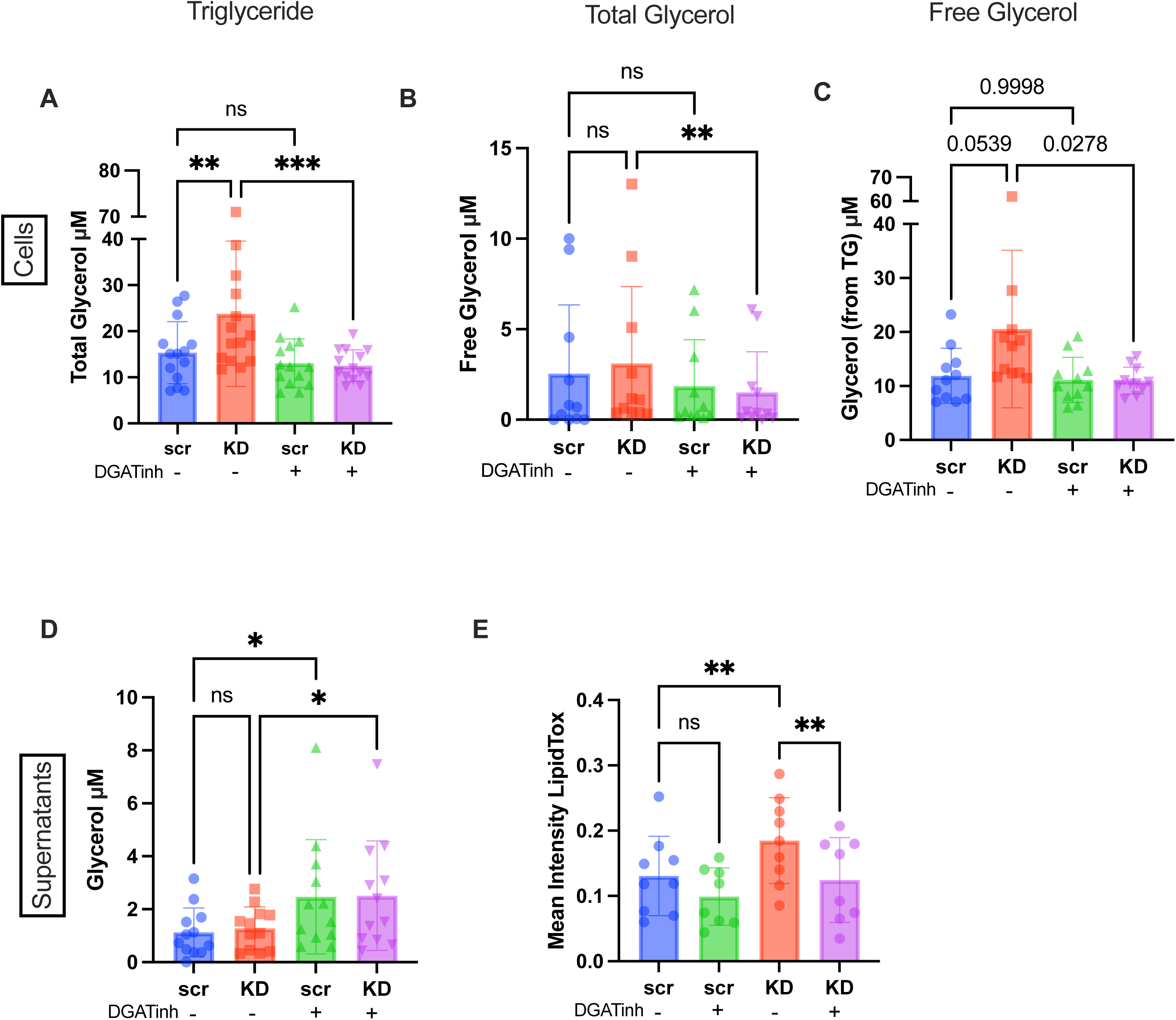
DGAT1 and DGAT2 inhibitors reduce lipid droplets in *TARDBP* knockdown MDMi. **A-C**) Total glycerol, free glycerol and triglyceride levels in cells, measured using Promega TriGlo Assay (N=14), with (+) and without (-) DGAT inhibitors (DGATinh), comparing scramble control (Scr) and *TARDBP* knockdown (TDP-43 KD). **D**) Total glycerol level in supernatants measured using Promega TriGlo Assay (N=12) +/- DGAT inhibitors (DGATinh), comparing scramble control (Scr) and *TARDBP* knockdown (TDP-43 KD). **E**) Mean Lipidtox intensity measured by confocal imaging and quantified by CellProfiler (N=9) in scramble control (Scr) and *TARDBP* knockdown (TDP-43 KD) +/- DGAT inhibitors (DGATinh). Statistical Analysis: 2-way ANOVA Sidak’s Multiple comparisons test Note: Free glycerol levels in supernatants are below kit detection. Total glycerol can be assumed to be triglycerides.

We then measured lipid droplet accumulation in the DGAT inhibitor-treated cells and found a significant decrease in the fluorescence intensity of lipid droplets in the *TARDBP* knockdown in treated compared with untreated cells; this effect was not seen in the scramble controls (**Fig. 6E**). Taken together, these data suggest that increased lipid droplet formation in the *TARDBP* knockdown is a result of TG accumulation, possibly a result of increased synthesis driven by DGAT1 and 2.

To further interrogate whether *TARBDP* knockdown MDMi are in fact accumulating triglycerides via increased synthesis, we examined the gene expression of DGAT1 and 2 in the knockdown and found, unexpectedly, that their expression was decreased **(Fig. S7A, B**). However, triglyceride hydrolysis genes ATGL and HSL were increased (**Fig. S7C, D**). This suggests that although the cells may not be increasing *de novo* triglyceride synthesis, triglyceride hydrolysis genes are still upregulated due to the increased accumulation. It is also possible that DGAT1 and 2 expression is reduced as a negative feedback mechanism to deal with increased enzyme activity. Additionally, DGAT2 is normally localized in the ER, but it is also found within lipid droplet membranes and facilitates lipid droplet expansion at the ER-lipid droplet interface. DGAT1, on the other hand, converts exogenous pre-formed fatty acids into triglycerides [64]. DGAT1 and 2 could therefore be involved in lipid droplet expansion and regulation of exogenous fatty acid uptake, thus resulting in decreased lipid droplet accumulation when they are inhibited.

### TDP-43 depleted MDMi have increased fatty acid uptake

We then wanted to examine whether *TARDBP* knockdown MDMi exhibit alterations in fatty acid uptake, which could explain the increase in TG synthesis or storage. *FABP4*, involved in regulating fatty acid uptake and transport, and *ELOVL3*, involved in fatty acid elongation, were among the most upregulated metabolism genes in the *TARDBP* knockdown. Additionally, *ACSL4* and *ACSL1* (Acyl-CoA Synthetase Long-Chain family), which encode enzymes that activate long-chain fatty acids by converting them into fatty acyl-CoA, were also upregulated. Based on this, we hypothesized that TDP-43 depletion causes increased fatty acid uptake, which could contribute to increased TG synthesis and lipid droplet accumulation. Indeed, we found a significant increase in the uptake of BODIPY-labeled C12 fatty acid in the *TARDBP* knockdown condition, as shown by quantification of confocal imaging (**Fig. 7A, B**).

**Fig 7:**
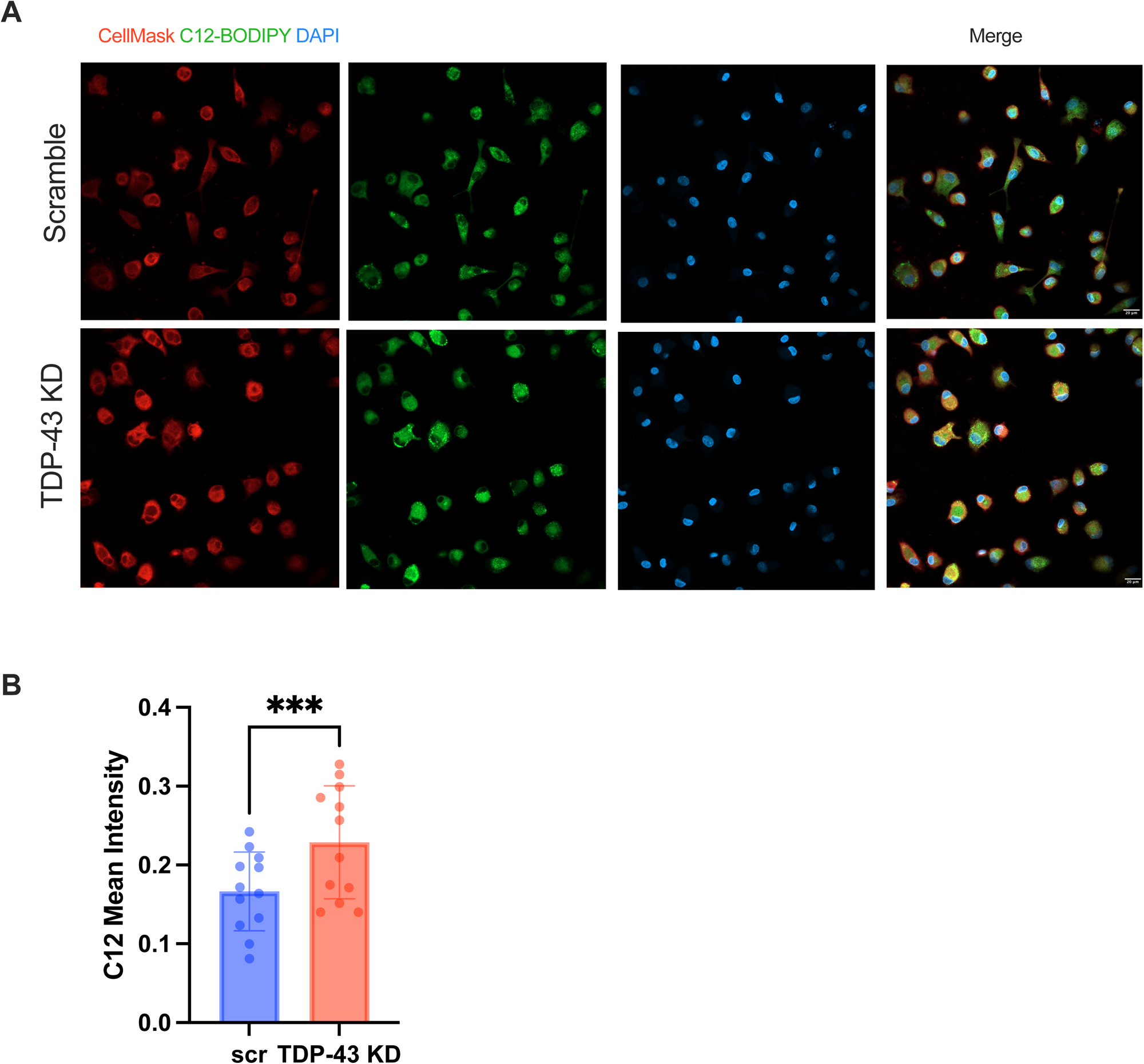
TDP-43 depleted MDMi show increased uptake of fatty acids. **A**) 20X Confocal images for BODIPY C12 uptake in scramble compared to *TARDBP* knockdown (TDP-43 KD), N=12. Scale bar = 20μm. **B**) Quantification of BODIPY C12 mean fluorescence intensity (MFI) calculated by CellProfiler. Statistical Analysis = paired t-test

### TDP-43 depleted MDMi have altered morphology and function

Finally, in addition to the effects on lipid metabolism, we wanted to understand the effect of *TARDBP* knockdown on conventional microglia functions. It is known that microglia cells change their morphology depending on their activation state and function. Surveilling and homeostatic microglia are thought to be ramified with more processes, whereas activated or phagocytic microglia have an ameboid morphology [65]. Here, we found that TDP-43 depleted MDMi had a more rounded morphology and a smaller cell body (**Fig. 8A**). This was quantified using the “compactness” metric in CellProfiler, where lower values indicate a more rounded morphology (**Fig. 8B**) [66]. Rounded microglia have been associated with enhanced phagocytosis [67], consistent with previous reports showing that microglia in ALS exhibit a more activated phenotype [4,35,68,69]. We then measured phagocytic capacity using a dextran uptake assay and found that TDP-43 depleted cells exhibited an increased average mean of dextran intensity, suggesting increased phagocytic activity (**Fig. 8C, D**).

**Fig 8:**
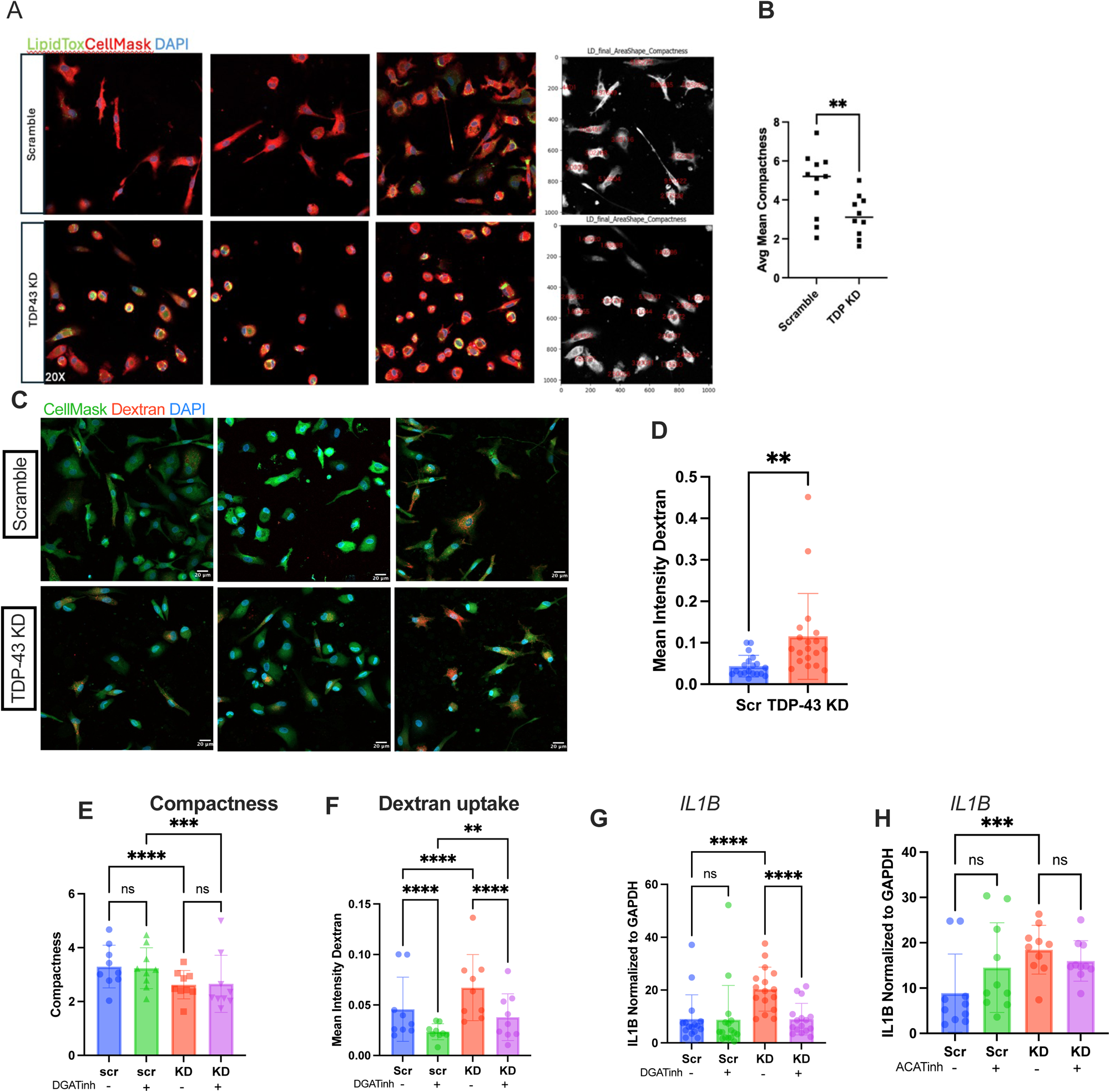
TDP-43 depleted MDMi have altered morphology and function. **A**) Confocal images for scramble control and *TARDBP* knockdown (TDP-43 KD) for MDMi from 3 individuals showing CellMask, LipidTox and DAPI stain. Panel 4 shows the compactness measure in CellProfiler using CellMask stain. **B**) Quantification of compactness measured by CellProfiler (N=10) in scramble and *TARDBP* knockdown (TDP KD). **C**) Confocal Images of Dextran uptake in scramble and *TARDBP* knockdown (TDP-43 KD), showing CellMask, Dextran and DAPI stain, 10X. Scale bar = 20μm. **D**) Quantification of Dextran uptake images using CellProfiler to determine mean dextran intensity within cells (N=10). **E**) Compactness measured by CellProfiler in scramble (Scr) and *TARDBP* knockdown (KD) +/- DGAT inhibitors (n=9). **F**) Dextran uptake quantification by CellProfiler in scramble (Scr) and *TARDBP* knockdown (KD) +/- DGAT inhibitors (N=9). **G**) qPCR for IL1B expression CellProfiler in scramble (Scr) and *TARDBP* knockdown (KD) +/- DGAT inhibitors (N=16). **H**) IL1B expression in scramble (Scr) and *TARDBP* knockdown (KD) MDMi treated with ACAT1 inhibitor. Statistical analysis for B&D: Paired t-test. Statistical Analysis for E-H: 2-Way Anova

### Phagocytosis and IL1***β*** levels are reduced in TDP-43 depleted MDMi treated with DGAT1 and DGAT2 inhibitors

To confirm that the inflammatory and phagocytic phenotypes observed in the *TARDBP* knockdown microglia are driven by elevated TG levels, we measured these outcomes in MDMi treated with DGAT inhibitors. We found no significant change in cell morphology (**Fig. 8E**). However, dextran uptake was reduced in DGAT inhibitor-treated MDMi (**Fig. 8F**), and *IL1*β expression was significantly reduced in the *TARDBP* knockdown but not in the scramble (**Fig. 8G**), which is in line with the effect on lipid droplets and triglyceride accumulation. We also found that ACAT1 inhibition did not alter *IL1*β expression (**Fig. 8H**), indicating that the *IL1*β gene expression increase is downstream of triglyceride accumulation. The fact that morphology was not altered upon DGAT inhibition suggests that it is not directly related to triglyceride or lipid droplet accumulation but may be a result of alterations in cytoskeletal proteins, which are known to be transcriptionally regulated by TDP-43 [70,71].

To understand how inhibiting triglyceride synthesis affects the pathways that were altered by *TADRBP* knockdown, we analyzed gene expression changes in DGAT inhibitor-treated samples with Fluidigm microfluidic qPCR. Interestingly, we found that most of the cholesterol biosynthesis genes remain downregulated with DGAT inhibition. However, key fatty acid genes and lipid droplet-associated genes like *FABP4*, *ELOVL3,* and *BSCL2,* which were upregulated in the knockdown, are reduced with DGAT inhibition. Many upregulated immune genes are also reduced significantly (**Fig. S8**). To determine whether DGAT inhibition reduces fatty acid uptake, we measured C12 uptake and found a significant reduction in the knockdown, but not in the scramble (**Fig. S9A**). C12 uptake can also be reduced by directly inhibiting a fatty acid receptor such as CD36 (**Fig. S9B**); however, we found that inhibition of CD36, using an inhibitor called SSO, did not result in decreased lipid droplet intensity (**Fig. S9C**). In fact, lipid droplet intensity was increased in SSO-treated control cells both with 4-hour and 24-hour treatment (**Fig. S9C, D**), suggesting that LD formation may not be primarily regulated by CD36-mediated fatty acid uptake, and involves more complex metabolic alterations.

### Lipidomic analysis of TARDBP knockdown MDMi demonstrates increases in monounsaturated triglycerides

To better understand the lipid profile in the TDP-43 depleted MDMi, we ran targeted LCMS-based lipidomic analysis and found alterations in many lipid species in the knockdown compared to the scramble control. Significant alterations were found in acylcarnitine (AC), triglyceride (TG), diacylglyceride (DG), and monoacylglyceride (MG) species (**Fig. 9A-D**). Notable trends in TG alterations included an increase in various mono and poly-unsaturated species of TGs (18:1 and 20:4), which are usually stored in neutral lipid droplets, while saturated TGs (18:0) were reduced (**Fig. 9E**), with TG 54:0/18:0 being significantly reduced in the knockdown (p<0.05). We also found alterations in Bis(monoacylglycero)phosphate (BMP), which are unique phospholipids found in the inner membranes of late endosomes and lysosomes and play an important role in lysosomal stability and lipid degradation (**Fig. S10**). Interestingly, polyunsaturated species were increased, whereas saturated species were reduced. Most species of mono-hexosyl ceramides (glucosylceramide), which are hydrolyzed in the lysosome to provide the cell with glucose and ceramide, were decreased, supporting the alterations in BMP that suggest defects in lysosomal degradation. Long-chain ACs (AC C12:0 and C18:0) were decreased while short-chain ACs (AC C3:0) were increased, which could indicate increased oxidation of fatty acids (**Fig. S10**). Overall, the lipidomics data suggests alterations in triglyceride and fatty acid metabolism, as well as possible dysfunction in lysosomal degradation.

**Fig 9:**
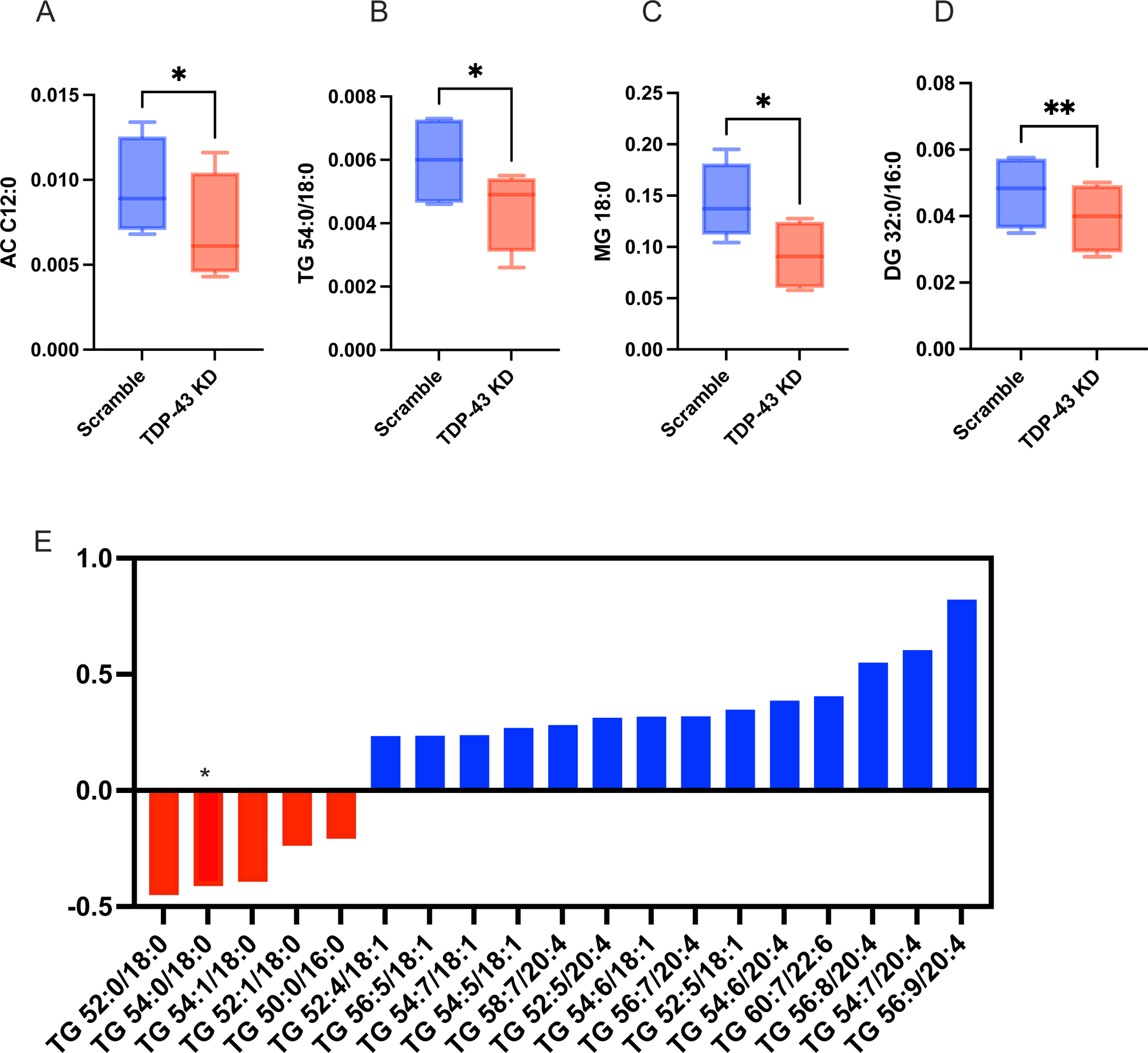
Lipidomic analysis of *TARDBP* knockdown MDMi demonstrates increases in monounsaturated Triglycerides. Box plot showing significantly altered lipid species in scramble vs *TARDBP* KD (TDP-43 KD): **A**) Acylcarnitine (AC C12:0) **B**) Triglyceride (TG 54:0/18:0) **C**) Monoacylglycerol (MG 18:0), and **D**) Diacylglycerol (DG 32:0/16:0). **E**) Bar chart showing fold change of triglyceride species in the knockdown compared to scramble control, with red bars showing decreased species and blue bars showing increased species. The asterisk shows significance (P<0.05). Statistical analysis: paired t-test, N=4

### ALS-patient derived MDMi show increased lipid droplets and IL1***β***, rescued by DGAT inhibition

We obtained peripheral blood mononuclear cells (PBMCs) from three individuals with mutations in *TARDBP*, one of whom was diagnosed with ALS and two of whom are pre-symptomatic mutation carriers, to make MDMi from (**Fig. S11A**). All three patients had missense mutations that have been shown to cause TDP-43 loss of function, mislocalisation, and aggregation [72–76], although *TARDBP* expression as measured by qPCR was unchanged (**Fig. S11B**). Compared to age and sex-matched controls, we found a significant increase in both *IL1*β gene expression and protein levels in the supernatants of the *TARDBP*-mutant (TDP-ALS) MDMi, which were reduced with DGAT inhibitor treatment (**Fig. 10A, B**). We also obtained PBMCs from patients with sporadic ALS (sALS) (**Supplementary Table 2**) and observed the same increase in *IL1*β expression and protein levels in their MDMi (**Fig. 10C-D**). Interestingly, lipid droplet intensity was increased in the TDP-ALS MDMi, but this only reached significance when including every image in the dataset rather than the mean of all images for each sample (**Fig. 10E and Fig. S11D**) and was rescued with DGAT inhibitor treatment (**Fig. 10E and Fig. S11C-D**). We also measured mean LD area and found that this was not significantly altered (**Fig. 10F**), although the maximum LD radius was significantly higher in the TDP-ALS MDMi (**Fig. 10G**), as seen in confocal images (**Fig. 10H**). DGAT inhibition did not have a significant effect on LD area or maximum radius in TDP-ALS MDMi (**Fig. 10F and Fig. S11E-F**), and in fact, DGAT inhibitor treatment seems to slightly increase LD area even though mean intensity was reduced. The sALS MDMi, on the other hand, showed a significant decrease in mean LipidTox intensity (**Fig. 10I**), whereas LD area and maximum radius were significantly increased (**Fig. 10J, K**), as also evident in imaging data (**Fig. 10L**). Again, significant trends were seen when every image was included in a nested analysis rather than using the mean of each donor (**Fig. S11H, I**), pointing to large variability between cells. This suggests that, while overall there were fewer LDs in sALS MDMi, resulting in lower mean intensity, there was a greater accumulation of large LDs.

**Fig 10:**
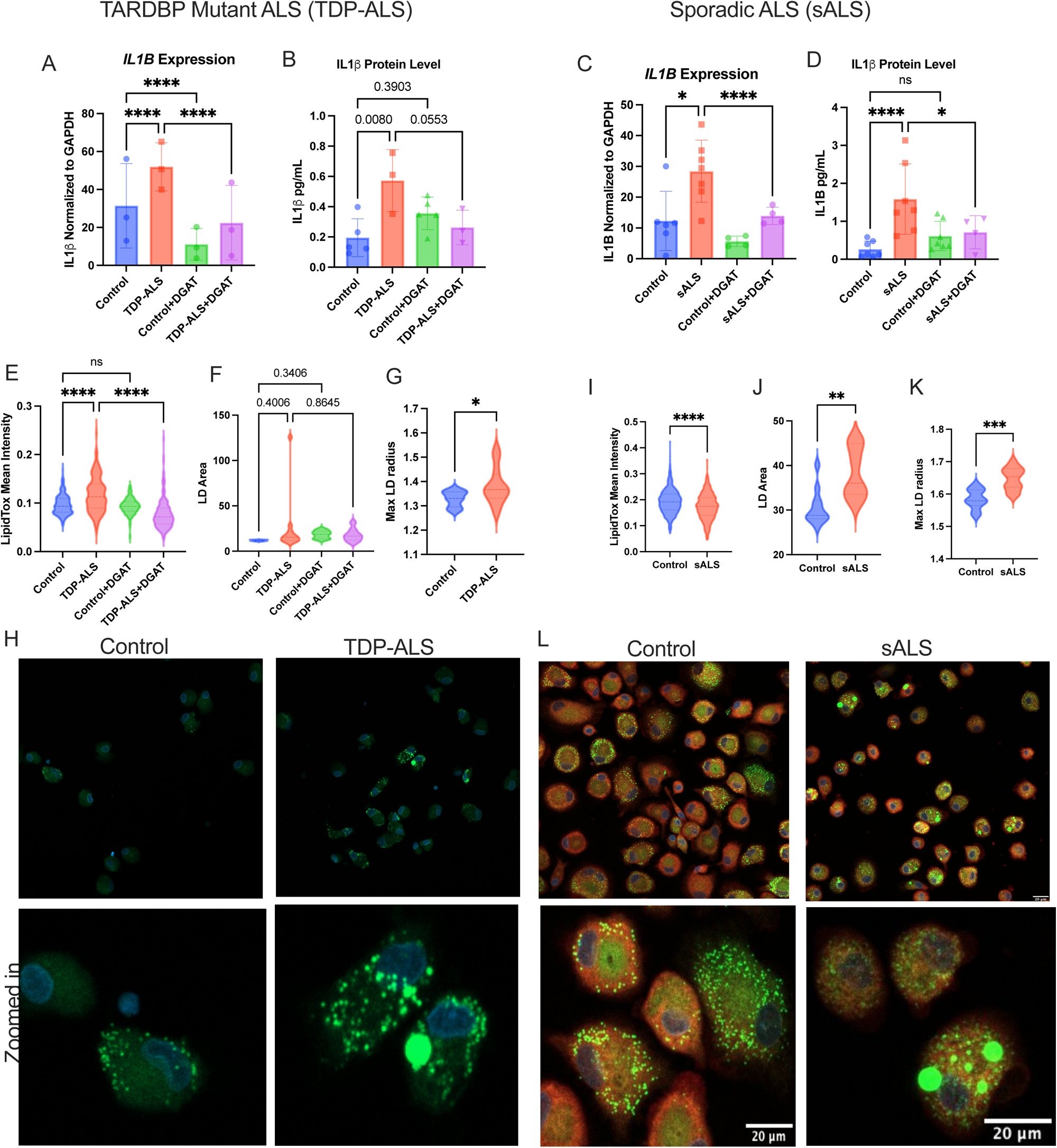
ALS-patient derived MDMi show increased lipid droplets and IL1β, rescued by DGAT inhibition. **A**) *IL1B* gene expression in TDP-ALS and matched control MDMi with DGAT inhibitor treatment using qPCR (N=3). **B**) IL1β protein quantification in supernatants of TDP-ALS and matched control MDMi using ELISA (N=3). **C**) *IL1B* gene expression in sporadic ALS MDMi and matched controls (N=7) with DGAT inhibitor (N=4). **D**) IL1β protein quantification in sporadic ALS and matched control MDMi supernatants (N=7). **E**) Quantification of mean LipidTox intensity for in TDP-ALS and control MDMi with DGAT inhibitor treatment (n=3). **F**) Quantification of mean lipid droplet (LD) area in control and TDP-ALS MDMi treated with DGAT inhibitors and **G**) Maximum lipid droplet (LD) radius using confocal images for LipidTox in control and TDP-ALS MDMi (N=3). **H**) Confocal images for control and TDP-ALS MDMi (20X) showing zoomed in images in the bottom panel with LipidTox and DAPI stain. **I**) Quantification of mean LipidTox intensity for control vs sporadic ALS MDMi (n=4). **J**) Quantification of mean lipid droplet (LD) area and **K**) Maximum lipid droplet (LD) radius using confocal images for LipidTox staining of control vs sporadic ALS MDMi (N=4). **L**) Confocal images for control and sporadic ALS MDMi (20X) showing zoomed in images in the bottom panel with LipidTox and DAPI stain. Scale bar = 20μm. Note: CellMask stain was not applied in the ALS-TDP samples and their corresponding controls. Statistical Analysis: A-D and F) 2-way ANOVA E) Nested 2-way ANOVA G-K) Nested Unpaired t-test.

Finally, we performed microfluidic qPCR analysis using Fluidigm on MDMi derived from TDP-ALS and sALS patients, along with healthy control samples and DGAT inhibitor-treated samples (**Fig. 11**). Notable differences were observed between TDP-ALS and sALS samples compared to healthy controls (**Fig. S12A**). Principal component analysis (PCA) revealed a clear separation between TDP-ALS MDMi and their age and sex-matched controls (**Fig. S12B**). In contrast, the separation between sALS samples and their matched controls was less distinct, suggesting greater heterogeneity or subtler transcriptional changes in the sALS group (**Fig. S12C**). Volcano plots show gene alterations in both TDP-ALS MDMi compared to matched controls, and sALS MDMi compared to matched controls (**Fig S12D-E**). There were not many genes significantly altered, even with unadjusted p-values, presumably due to the low sample numbers and individual heterogeneity.

**Fig 11:**
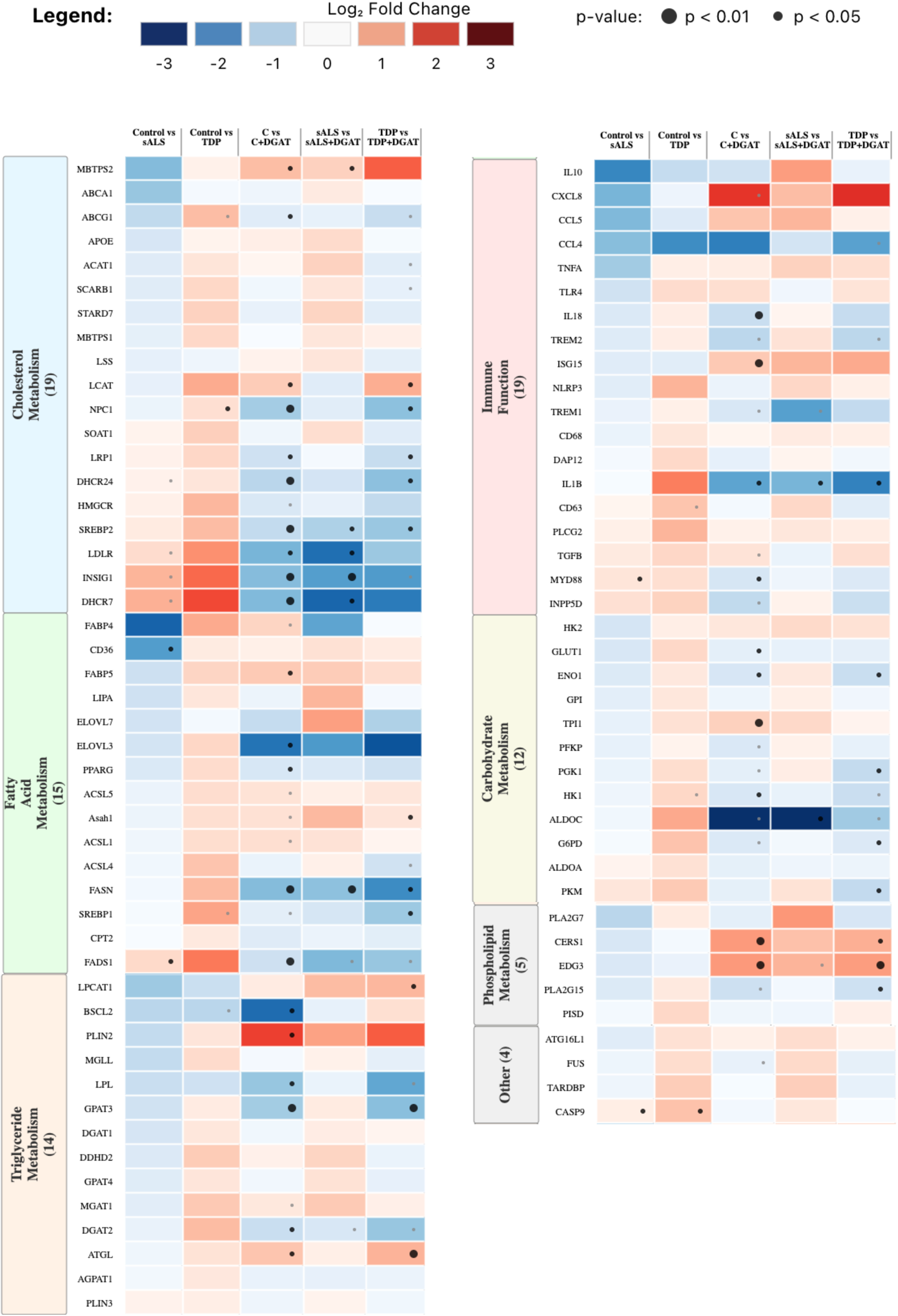
Fluidigm analysis of ALS patient-derived MDMi shows distinct alterations. Heatmaps showing Fluidigm data for gene expression alterations in control versus sporadic ALS (sALS) and control versus TDP-mutant ALS (TDP) groups, along with DGAT inhibitor-treated groups. Fold change is expressed as Log2FC, and values that were significant are indicated by the large black dot (p<0.01) and small black dot (p<0.05), while the grey dots indicate close to significance (p=0.05). Statistical analysis for control versus sALS and control versus TDP: multiple unpaired t-test. Statistical analysis for DGAT-treated versus untreated: multiple paired t-test.

We also found divergent gene expression patterns between the patient-derived MDMi and shRNA knockdown MDMi. The TDP-ALS MDMi showed increased expression of many cholesterol biosynthesis genes (*DHCR7*, *INSIG1*, *SREBP2*), which were all reduced with DGAT inhibitors (**Fig. 11**). This contrasts with the knockdown, where *SREBP2* and *INSIG1* were significantly downregulated (**Fig. 2D**). Some consistencies with the knockdown were the increased expression of triglyceride and fatty acid metabolism genes in the TDP-ALS MDMi, which are not seen in the sALS MDMi (including *FABP4*, *FADS1*, *MGAT1,* and *DGAT2*) (**Fig. 11**). These data suggest that altered fatty acid and triglyceride metabolism may be specific to mutations in *TARDBP*, and possibly nuclear depletion of TDP-43, whereas in sALS cases, there may be other factors contributing to disrupted lipid metabolism that cause LD accumulation and altered immune responses.

Taken together, our data demonstrate that TDP-43 plays a critical role in regulating lipid metabolism in microglia. Specifically, nuclear depletion of TDP-43 results in an accumulation of triglycerides, likely via increased uptake of fatty acids and upregulated triglyceride synthesis caused by bioenergetic alterations. TDP-43 nuclear depletion also results in increased phagocytic capacity of microglia and higher secretion of pro-inflammatory cytokines like IL1β, which is downstream of triglyceride accumulation. Patients with mutations in *TARDBP* exhibit similar phenotypes, which are reversed with DGAT inhibition, suggesting that lipid pathways, specifically triglyceride metabolism may be dysregulated in *TARDBP*-driven ALS. MDMi from sALS patients also accumulate large LDs, but show distinct gene expression profiles, suggesting a different mechanism of LD accumulation.

## Discussion

Although mutations in *TARDBP* are rare and make up only 5% of the total familial ALS population, TDP-43 pathology is present in over 90% of ALS patients. Additionally, it is observed in over 50% of Alzheimer’s disease (AD) cases and some Parkinson’s disease (PD) cases [77–81]. Rather than focusing on specific TARDBP mutations, we employed a general loss-of-function model to investigate the broader cellular consequences of TDP-43 dysfunction, aiming to identify mechanisms relevant across multiple neurodegenerative conditions. Loss-of-function models effectively mimic the pathological effects of TDP-43 aggregation [82–84], and in fact, recent studies have shown that inducing TDP-43 aggregation leads to loss of nuclear TDP-43 [85,86]. Notably, TDP-43 depletion from microglia has been shown to exacerbate neuroinflammation, pointing to the importance of studying TDP-43 pathology in non-neuronal cells [87].

In our study, we achieved significant nuclear depletion of TDP-43 using shRNA knockdown in monocyte-derived microglia-like cells (MDMi). Unlike patient-derived MDMi in previous studies, our model did not exhibit cytoplasmic aggregation or phospho-TDP-43, suggesting that nuclear depletion alone is insufficient for these pathological features but still drives significant metabolic and immune alterations. Our data demonstrates a clear link between TDP-43 nuclear depletion and triglyceride alterations that contribute to lipid droplet (LD) accumulation and altered immune responses in MDMi. While previous studies have demonstrated that microglia LD accumulation can occur via upregulation of triglyceride pathways [63], this has not been shown in the context of ALS. Excessive LD accumulation could be a consequence of several factors, and their varying composition can affect functional outcomes. In a study using both human AD brain tissue and iPSC-derived microglia with different ApoE genotypes (ApoE3/E3 vs ApoE4/E4), microglia with ApoE4/E4 genotype had increased lipid droplets, greater expression of ACSL1 when exposed to fibrillar Aβ and an induction of triglyceride synthesis. These cells were also found to express markers of “cellular senescence” and exhibit less phagocytosis [25]. Here, we found that triglyceride-driven LDs in fact result in increased phagocytic activity, which points to heterogeneity in the effects of LD accumulation.

Several genes associated with fatty acid metabolism, including *ACSL1*, *FABP4, ELOVL3, and ACSL4,* were upregulated in our knockdown. In addition to the involvement of *ACSL1* with LD-accumulating microglia [25,88], some of these other genes have also been linked to lipid dysregulation and ALS. IPSC-derived microglia with mutations in profilin-1(PFN1) that are causative for ALS were shown to exhibit upregulated *FABP4* and *FABP5* [89]. Other studies have linked *FABP4* to increased triglyceride synthesis and lipid droplet accumulation [40]. *ELOVL3* was found to be significantly increased in *LPL* knockdown microglia, which also exhibit increased LD formation, reduced cholesterol synthesis and efflux, and increased inflammation [44], which is in line with our measured phenotypes. In our knockdown, these genes were downregulated with DGAT 1 and 2 inhibitors, suggesting that inhibiting triglyceride synthesis triggers a feedback mechanism that downregulates fatty acid uptake/metabolism pathways.

Metabolic alterations and bioenergetic shifts are well-documented in ALS, with many studies reporting a transition from glycolysis to fatty acid oxidation, particularly in glycolytic tissues such as skeletal muscle [90–92]. Recent findings from our group [93] demonstrated that motor neurons from ALS SOD1 mutant mice exhibit upregulation of glucose, fatty acid, and amino acid catabolism, with impaired oxidative phosphorylation (OXPHOS) and increased fatty acid oxidation. In our current study, we found significant downregulation in the expression of *PFKP* and *G6PD*, both critical for glycolysis. Prior studies have linked TDP-43 to PFKP regulation, with TDP-43 loss-of-function reducing *PFKP* expression and activity, potentially through cryptic exon inclusion [50,94–96]. Our findings align with these reports, supporting a model in which loss of nuclear TDP-43 leads to impaired glycolysis and a shift toward fatty acid uptake and utilization.

*TARDBP* knockdown resulted in notable changes in immune-related gene expression. *TREM2* and *IL10* were significantly downregulated, while *TREM1*, *IL1*Β, and *CCL4* were upregulated. The opposing roles of TREM1 and TREM2 in inflammation suggest that loss of nuclear TDP-43 skews microglia toward a pro-inflammatory state [16,97,98]. Interestingly, *TREM2* is upregulated in DAM and microglia associated with neurodegeneration, but not in LDAM (lipid droplet-accumulating microglia), which are thought to be associated with aging and Alzheimer’s Disease [24]. Although sequencing studies have shown an upregulation of TREM2 (along with other DAM genes) in ALS microglia, it is known that TREM2 genetic variation (such as in the R47H variant) is also associated with ALS. Additionally, TREM2 is required for the protective role in attenuating the expression of pro-inflammatory mediators, including iNOS, TNFα, IL-1β, and IL-6, as well as mediating phagocytosis of TDP-43 aggregates in the context of ALS. Our data demonstrating a decrease in soluble TREM2 protein therefore supports defects in immune responses in *TARDBP* knockdown MDMi. The observed increase in expression of *CCL4*, which encodes a chemokine that correlates positively with better ALS functional scores [99], suggests a possible early-stage protective response that could become detrimental over time. Notably, treatment with DGAT inhibitors reduced *IL1*Β, *CCL4*, and *TREM1* expression, but did not alter *TREM2*, linking triglyceride accumulation specifically to pro-inflammatory pathways (21,93). Treatment with DGAT inhibitors also significantly reduced *NLRP3, IL1*Β, and *IL18* gene expression, and more importantly, IL-1β protein levels, implicating the inflammasome in the increased inflammatory phenotypes observed in the *TARDBP* knockdown.

The role of lipid droplet composition in microglia function remains poorly understood. While TREM2-deficient microglia accumulate cholesterol ester-rich LDs, which can be rescued by ACAT1 inhibitors [100], our data suggest that TDP-43 deficiency leads to triglyceride-driven LD accumulation, which cannot be rescued by ACAT1 inhibitors. Although we do see a decrease in soluble TREM2 protein levels in our model, our LD phenotype is driven by triglycerides rather than cholesterol esters. Notably, we found that inhibiting ACAT1 reduced LDs in the scramble control, whereas inhibiting DGAT1 and 2 reduced LDs in the knockdown, suggesting that TDP-43 depletion leads to a shift in LD composition in MDMi. Here we observe a distinct phenotype where LD accumulation is accompanied by increased phagocytosis, unlike other studies that have shown impaired phagocytosis in LD-accumulating microglia [24,25,100,101]. Some studies have also demonstrated the importance of LD accumulation in driving anti-inflammatory responses in microglia [102,103]. This highlights the fact that heterogeneity in LD-associated phenotypes needs further characterization in the context of neurodegenerative diseases involving lipid alterations. In our model, nuclear depletion of TDP-43 could be driving triglyceride-mediated inflammatory pathways, which increase baseline activation of MDMi and increase phagocytic activity.

Our lipidomic analysis supports the observed phenotypes, as we found several unsaturated TGs to be elevated in the *TARDBP* knockdown. These were primarily TGs containing oleic acid (18:1), which are known to accumulate in neutral lipid droplets. Interestingly, previous studies have shown accumulation of oleic acid (OA18:1) in a model of Parkinson’s Disease [104,105], and macrophages are known to accumulate unsaturated TGs in the “M1 polarization state”, associated with inflammation [106]. Acyl carnitines (ACs) are fatty acids that are transported to mitochondria for beta-oxidation, and alterations are an indication of mitochondrial dysfunction. The decrease in long-chain ACs (C12:0) and corresponding increase in short-chain ACs (AC 3:0) suggests increased fatty acid oxidation. This could also potentially implicate incomplete fatty acid oxidation as a consequence of excessive fatty acid uptake for ATP production in a state of impaired glycolysis (supported by the downregulation of glycolytic genes in our Fluidigm data). Accumulation of short-chain ACs is also associated with inflammatory responses and oxidative stress in a variety of metabolic contexts [107], and alterations in short-chain ACs have been found in neurodegeneration and aging [108,109]. Finally, we found reductions in BMPs and lactosylceramides, and an increase in mono hexosyl ceramides, both of which are an indication of impairments in lysosomal function and lipophagy [110–112]. This could imply an alternate mechanism of triglyceride accumulation that results from defects in lysosomal degradation of lipids, rather than increased *de novo* synthesis.

Overall, our findings suggest a *TARDBP*-specific metabolic shift, characterized by reduced cholesterol biosynthesis and uptake, coupled with a possible impairment in glycolysis, resulting in increased fatty acid uptake and triglyceride accumulation. The findings of triglyceride alterations are not unique to our study and have been observed in ALS patient serum, Parkinson’s Disease iPSC-motor neurons, as well as ApoE4 microglia [63,113–116], although AD lipidomic studies have found more drastic alterations in phospholipids, sphingolipids, and cholesterol esters [117–120]. However as described above, the LD phenotype observed in our TDP-43-depleted MDMi is unique in its effect on microglia function, and this could be explained by bioenergetic alterations that specifically affect glycolysis and fatty acid metabolism in these cells. Interestingly, a previous study using *TARDBP* knockdown in iPSC-derived motor neurons and HeLa cells found reduced ATP-linked respiration and impaired mitochondrial function, reduced *PFKP* expression, and no alteration in triglycerides [50]. While our data also show reduced *PFKP* expression, the increase in triglycerides appears microglia-specific, suggesting unique metabolic adaptations across cell types. Additional studies examining substrate utilization in *TARDBP* knockdown microglia and patient-derived microglia will be crucial in elucidating the mechanisms underlying lipid droplet heterogeneity.

Lastly, we were able to show that MDMi derived from individuals with *TARDBP* mutations recapitulated some key phenotypes of our *TARDBP* knockdown MDMi, including LD accumulation, increased IL1β, and response to DGAT inhibitors. This provides an exciting, new understanding of metabolic alterations that could drive ALS pathology in a TDP-43-dependent manner. MDMi derived from sALS patients also displayed significant increases in IL1β and accumulation of large LDs, although mean LD intensity was reduced. 90% of sALS patients exhibit TDP-43 pathology, and these phenotypes could therefore be driven by TDP-43 to some extent. However, gene expression analysis showed higher expression of fatty acid and triglyceride metabolism genes in the TDP-ALS MDMi compared to sALS MDMi, specifically, *DGAT2* was increased in the TDP-ALS MDMi and not sALS MDMi (in addition to *FABP4*, *FADS1*, *FASN,* and *ACSL4*) although only *FASN* was significant (p-value<0.05). Measuring TDP-43 nuclear depletion by immunohistochemistry in TDP-ALS compared with sALS MDMi would enable a better understanding of whether the differences are driven by TDP-43 nuclear depletion. It is important to note that although DGAT inhibitors reduced LipidTox intensity, they did not reduce LD size in TDP-ALS MDMi, and in fact increased LD maximum radius, suggesting that while blocking DGAT enzymes may reduce the number of LDs, the existing triglycerides may be stored in fewer LDs, making them bigger. Additionally, DGAT1 is thought to be involved in the formation of new LDs, while DGAT2 is involved in expansion of existing LDs [64,121]. The inhibitors may have a disproportionate effect on these enzymes, causing greater inhibition of DGAT1 compared to DGAT2.

Interestingly, other immune-related genes like *TREM2* and *CCL4* showed distinct alterations in the TDP-ALS MDMi compared to the knockdown. *TREM2* was slightly upregulated, while *CCL4* was downregulated. This could be a difference in the stage of disease captured by both models or could reflect heterogeneity in ALS subtypes. When comparing to the shRNA knockdown, genes related to carbohydrate metabolism and cholesterol biosynthesis were both increased in the TDP-ALS MDMi, which contrasts with the *TARDBP* knockdown MDMi. This could be a compensatory effect of reduced function or suggest alternate mechanisms of lipid dysfunction in both models. Functional assays to measure glycolysis and cholesterol synthesis would therefore be required to elucidate how the knockdown model differs from patient-derived MDMi and the extent to which gene expression may be correlated with phenotypes such as triglyceride and cholesterol levels in cells.

Our study has certain limitations to be acknowledged. While several methods exist to model human microglia, we used monocyte-derived microglia-like cells (MDMi), which undergo a similar polarization step as iPSC-derived microglia, but are more adept at incorporating human variability driven by age, disease state, and natural heterogeneity [122,123]. Although this heterogeneity reflects real-world diversity, future studies should employ a larger sample size to determine whether there are genotype-specific effects. Further, all our analyses were performed at Day 14 post-differentiation, which may represent a late-stage response to *TARDBP* knockdown since the shRNA treatment is done on Day 4. Additionally, the observed increase in *ATGL* and *HSL* expression and decrease in *DGAT1* and *DGAT2* expression in the knockdown model, despite elevated triglycerides, suggests that microglia may attempt to counteract triglyceride accumulation over time. Time-course experiments could clarify whether early-stage metabolic adaptations differ from later responses.

Our targeted gene expression approach with Fluidigm analysis, while informative, does not capture the full transcriptomic landscape. Unbiased RNA sequencing could provide a more comprehensive view of *TARDBP* knockdown effects and uncover important differences between the shRNA knockdown model and the ALS patient-derived MDMi model. Additionally, lipidomic analysis of isolated lipid droplets from a greater sample size to understand the full profile of triglyceride, cholesterol and fatty acid species would help validate some of our data. With regards to the use of shRNA-mediated lentiviral knockdown, while this is effective and provides a very stable knockdown allowing for downstream assays, alternative approaches such as siRNA or CRISPR-mediated knockdown in iPSC-derived microglia could further validate our findings. Our knockdown reduced protein levels by 20-30%, which is low compared to other studies that use iPSC-derived microglia or cell lines that allow for more efficient knockdown, however, even with a modest knockdown, we do see a large biological effect. Nevertheless, using these alternate approaches would also provide larger cell counts to perform activity assays and western blots to confirm gene expression data. Lastly, while we were able to obtain three TDP-ALS and nine sALS patient samples, cell yields were insufficient to perform nuclear TDP-43 staining or functional assays such as phagocytosis and response to LPS stimulation. Future studies with a larger cohort of patient-derived samples will be essential to fully elucidate how lipid pathways are altered in ALS and how these changes impact immune function. RNA sequencing and splicing analysis would also potentially uncover novel targets by which TDP-43 could directly modulate bioenergetic and lipid pathways in microglia.

## Conclusion

Our study demonstrates that nuclear depletion of TDP-43 in MDMi leads to metabolic and immune alterations, characterized by increased fatty acid uptake, triglyceride accumulation, and possible impairment in glycolysis. These changes are accompanied by pro-inflammatory cytokine production, independent of cytoplasmic TDP-43 aggregation. The findings highlight the importance of microglial lipid metabolism in neuroinflammation and suggest that triglyceride accumulation may drive microglia activation in ALS. Further studies on lipid droplet heterogeneity and metabolic adaptations in microglia could provide new insights into ALS pathogenesis and identify potential therapeutic targets.

## Materials and Methods

### MDMi Cell Culture

MDMi are created as described previously [124]. Blood from healthy human donors is separated using a density gradient medium Lymphoprep (Stemcell technologies #07851) to isolate mononuclear cells. These peripheral blood mononuclear cells (PBMCs) are cryopreserved in Fetal Bovine Serum (FBS) with 10% Dimethyl sulfoxide (DMSO) at -80C until needed. PBMCs are thawed and monocytes are obtained through CD14+ microbead isolation (Miltenyi #130-050-201). Monocytes are then plated in 96 well plates at a density of 200,000 cells per well and cultured in serum-free RPMI (Gibco #R8758) media with 1% penicillin and streptomycin (10,000 U/mL) (Fisher Scientific 15-140-122) and 2.5 μg/mL Fungizone (Cytiva #SV30078.01). A cytokine cocktail consisting of macrophage colony-stimulating factor (M-CSF) (10 ng/mL), granulocyte-macrophage colony-stimulating factor (GM-CSF) (10 ng/mL), nerve-growth factor-b (NGF-b) (10 ng/mL), chemokine ligand 2 (CCL2) (100 ng/mL), and interleukin-34 (IL-34) (100 ng/mL), is added to the media. Cells are differentiated into microglia-like cells (MDMi) over 10 days with the help of these cytokines. Cytokines were purchased from R&D Systems (NGF-b, GM-CSF, and IL-34) and Biolegend (M-CSF and CCL2). Monocyte-derived microglia-like cell models have been reviewed and characterized as an appropriate model to study human microglia *in vitro* [35,124–127]. Assays were done using MDMi from 3-6 individuals per experimental run. For each assay, 2 to 4 batches (repeat experimental runs) were done and the total number of individuals used per assay is in the figure legend.

### Preparation of shRNA lentiviral particle

Lentiviral particles were prepared as previously described [128,129]. Briefly, on day 1, 293T cells were transfected using Lipofectamine 2000 (Thermo Fisher Scientific, Waltham, MA, United States) with packaging and envelope plasmids (Vpx cDNA and pHEF-VSVG). On day 2, 293T culture media was replaced with RPMI-1640 Glutamax (Invitrogen, Waltham, MA, United States) containing 1% fungizone (Amphotericin B) and 1% penicillin/streptomycin. After 48 h, lentiviruses containing the Vpx particles were harvested, centrifuged for 5 min at 400 g and the supernatant collected. The supernatant was filtered using a 0.45-μm syringe filter (EMD Millipore, Burlington, MA, United States). Lentiviral particles containing targeted shRNA for each gene were obtained from Milipore Sigma (TARDBP construct: TRCN0000016038, Target Sequence: GCTCTAATTCTGGTGCAGCAA).

### Lentiviral mediated knockdown of MDMi

For the transduction of MDMi cells, on day 4 of differentiation, the culture media was replaced with 100 μl of Vpx-VLP and 100μL of fresh RPMI media containing 2X concentration of cytokines. After 2-3 hours, 10 μl TRC virus-containing shRNA or scramble control (Sigma) was added to each well. On day 7, puromycin (Life Technologies, Carlsbad, CA, United States) at a concentration of 3 μg/ml was added to eliminate non-transduced cells. On day 10, MDMi were lysed for RNA isolation [128,129]. A > 50% knockdown of RNA (by qPCR) was considered optimal for the experiment (higher knockdown efficiencies could not be obtained for the *TARDBP* gene, and this level of knockdown showed significant protein level reduction, so it was considered sufficient to study effects of *TARDBP* reduction). The knockdown was performed with PBMCs from 20 individuals (11 male, 7 female, and two unknown, aged 18 to 70). Of these, 5 samples were removed from analysis due to insufficient knockdown (<50%), leaving 6 males, 7 females and 2 unknown. Information on the age, sex and ethnicity of these individuals are provided in **Supplementary Table1**.

### Cholesterol uptake, fatty acid uptake and lipid droplet staining

A fluorescent cholesterol analog, NBD-cholesterol (22-(N- (7-Nitrobenz-2-oxa-1,3-Diazol- 4- yl)Amino)-23,24-Bisnor-5-Cholen-3β-OI) (ThermoFisher N1148) was used to determine cholesterol uptake. Cells were incubated with NBD-cholesterol for 1 hour, followed by fixing with PFA and confocal imaging. Fatty acid uptake was measured by incubating cells in a similar manner with 20 μM C-12 BODIPY labeled fatty acid (ThermoFisher D3823). Staining of lipid droplets was performed using HCS LipidTox™ Deep Green or Red neutral lipid stain (ThermoFisher H34475) according to manufacturer instructions. Confocal imaging was used to obtain images. All images were quantified using CellProfiler.

### Total Cholesterol/Free cholesterol Assay

Total and free cholesterol levels were determined using the Amplex Red cholesterol assay (ThermoFisher A12216) as per the manufacturers protocol. Cells were grown on 12 well plates, and after the knockdown protocol on day 10, media containing viral particles was replaced with fresh RPMI media without cytokines. After 3 days, cells were collected using a cell scraper, and frozen at -80°C in 250 μL water. Supernatants were also collected and frozen. When ready to perform the assay, cells were thawed, and the protein was quantified using Bradford’s assay. Lipid extraction was done by adding chilled chloroform and methanol in a 2:1 ratio. This was followed by vortexing for 30 seconds and 2-minute incubation on ice, repeated 3 times. The samples were then centrifuged at 14000rpm for 10 minutes, followed by transferring the lower organic layer to a new Eppendorf tube. This was placed under nitrogen until all the chloroform was dried, followed by resuspension of the lipids in the reaction buffer provided in the kit. Cholesterol esters were calculated by subtracting free cholesterol from total cholesterol. All values were normalized to protein levels.

### Triglyceride/Glycerol Assay

Triglyceride-Glo Assay (Promega J3160) was used to measure total glycerol and free glycerol in cells and supernatants, as per the manufacturer’s instructions. Cells were seeded in 96-well plates in duplicate (to perform the assay with and without lipase). On Day 10, media containing viral particles was replaced with fresh RPMI media without cytokines.

After 3 days incubation in fresh RPMI, on Day 14, the supernatant was removed and collected, and cells were washed once with PBS, followed by the addition of the glycerol lysis buffer (as per kit instructions). Cells and supernatants were assayed at the same time, and luminescence was measured by a plate reader. Triglyceride levels were calculated by subtracting free glycerol from total glycerol. An equal number of cells plated per well serves as normalization, as protein quantification could not be done with this kit.

### Immunohistochemistry

MDMi were plated in 24-well plates with glass coverslips. On Day 10, cells were washed 3X with 3% BSA and 0.1% TritonX in PBS, fixed with 4% PFA for 15 minutes, washed again, and incubated with primary antibodies (Goat Anti-Iba1, Fujifilm CAT# 011-27991 and Rabbit Anti-TDP43, R&D Systems, CAT# MAB7778) overnight at 4C. The next day cells were washed 3X with PBS followed by incubation with the appropriate Alexa-conjugated secondary antibodies for 1 hour at RT. Cells were washed again 3X, and the glass coverslips were then mounted on microscope slides using FluoroG mounting solution and imaged at 60X using confocal microscopy.

### Western Blot Analysis

Cell protein concentrations were measured using the Quick Start Bradford Protein Assay Kit 1 (Bio-Rad 5000201) in a Tecan Infinite F200 PRO spectrophotometer. 10 μg protein was combined with 4X Laemmli loading buffer in a final volume of 30 μL, heated at 95C and loaded on an Invitrogen 4–20% tris–glycine SDS-PAGE gel. Electrophoresis was conducted at 80-120 V using standard tris–glycine running buffer. The sample was transferred to an Immuno-Blot PVDF membrane (Bio-Rad 1620177) in standard tris–glycine transfer buffer with 20% methanol and 0.04% SDS at 150 mA for 2 h in a wet transfer. The primary antibodies used are as follows: Anti-TDP43, R&D systems, CAT# MAB7778, Anti-GAPDH Cell Signaling, CAT#5174S.

### Dextran Uptake Assay

Cells were incubated with 0.1 mg/mL Dextran Alexa Fluor™ 647 10,000 MW (ThermoFisher D22914) for 1 hour at 37C, followed by fixation with 4% PFA and imaging with a confocal microscope. As a positive control, cells were incubated with 10µM Cytochalasin D (FisherSci., Cat# 12-331) for 20 minutes prior to incubation with Dextran. Image analysis was done using CellProfiler.

### Cell Lysis and RNA Isolation

MDMi are lysed with RLT buffer (Qiagen #74104) with 1:100 β-mercaptoethanol, purified RNA is extracted using the RNeasy 96 well plate isolation kit (Qiagen #74182)._Reverse Transcriptase PCR and qPCR: RNA is transformed to cDNA using reverse transcription PCR. The PCR mix consists of dNTP mix (#R72501), Random Hexamers (#N8080127), RNase Inhibitor (#N8080119), MgCl2 (#AB0359), 10x PCR buffer (#4486220), and M-MLV Reverse Transcriptase (#28025013) purchased from Thermofisher Scientific. Reagents and RNA are loaded into a 96-well plate to a total volume of 50µL and run in an Applied Biosystems MiniAmp Thermocycler. Thermocycler program: 25°C for 10 minutes, 48°C for 45 minutes, 95°C for 5 minutes, and held at 4°C upon run completion.

Sample cDNA is loaded into a 96-well PCR plate with Taqman Fast Advanced Mastermix (#4444554), assay primer for the target gene to be detected with FAM, and housekeeping gene to be detected with VIC. qPCR primers were purchased from Thermofisher Scientific. For all experiments, the housekeeping gene glyceraldehyde-3-phosphate dehydrogenase (GAPDH) was used to normalize all values as relative expression (R.E.). Samples are run with the protocol: Stage 1 (x1): 50°C for 2 minutes, 95°C for 2 seconds. Stage 2 (40x): 95°C for 1 second, 60°C for 20 seconds. Cycle threshold (Ct) values are collected and normalized to GAPDH Ct.

### Microfluidic qPCR Analysis

Gene expression analysis was performed by parallel qPCR using the high-throughput Fluidigm BioMark HD platform (Standard BioTools, San Francisco, CA, USA), according to the manufacturer’s instructions. GAPDH and TUBB were used as reference genes, but final data are reported using GAPDH as a reference. A pre-amplification step was included to increase the number of cDNA copies to a detectable level and to allow the concurrent amplification of the different gene expression targets. The Fluidigm IFC was primed with control line fluid on the IFC controller. Subsequently, assay and sample mixes were loaded on the IFC and placed on the controller which pressure-loaded the assay components into the reaction chambers. The IFC was then placed on the Biomark HD for thermocycling and fluorescence detection. The data was reviewed on the Fluidigm Real Time analysis software. Data from multiple Fluidigm runs was normalized by calculating fold change of the knockdown over scramble control. Multiple paired t-test was used to determine significance of alterations, and this was used to make a volcano plot. Although Fluidigm analysis can only analyze 96 genes simultaneously, our data includes 110 genes, as multiple Fluidigm runs were combined, and not all genes were run on every sample.

### Confocal Microscopy

Cells were imaged on the Zeiss LSM 900 confocal microscope. Images were processed using FIJI image processing software and analyzed using the open-source program CellProfiler to classify and count cells and measure staining intensity. For CellProfiler, customized pipelines were developed to analyze lipid droplet intensity, TDP-43 localization and dextran uptake [66]. Compactness was measured using the “MeasureObjectShapeSize” module, where compactness is calculated as “the mean squared distance of the object’s pixels from the centroid divided by the area”. A filled circle will have a compactness of one, whereas flatter or more elongated and ramified cells will have a compactness greater than one.

### CellProfiler Pipelines

Imaging analysis pipelines used involved identifying cells as a primary object using DAPI stain, setting appropriate thresholds for the nucleus size. CellMask staining was used to identify the cell body as a secondary object. Cells were filtered to only include those with a nucleus (using “FilterObject” and “RelateObject” modules), followed by measuring the average intensity per cell (using “MeasureObjectIntensity” for LipidTox staining, NBD-cholesterol, or Dextrain staining. To measure TDP-43 nuclear and cytoplasmic staining, Hoechst staining was used to identify the nucleus, and Iba1 staining was used to identify the cell body. The “MaskObject” module was used to identify cytoplasmic area (without nucleus). Mean values per image were used for final analysis in all imaging analysis. At least 5 images per well were analyzed for each experiment.

### CellMask Stain

For lipid droplet staining and dextran uptake assays, CellMask (Invitrogen CAT#C10045) was used to stain the plasma membrane before fixing cells. Briefly, cells were incubated with CellMask (1:1000) for 5-10 minutes at 37°C in RPMI media, followed by washing 3X with PBS and fixing with PFA.

### Drug Inhibitors

DGAT 1 and 2 inhibitors (Sigma PF04620110 and PF06424439 respectively) were added simultaneously at a concentration of 5uM. To inhibit cholesterol esterification, 20 μM 3-[Decyldimethylsilyl]-N-[2-(4-methylphenyl)-1-phenethyl] propanamide (Sandoz 58-035), a specific ACAT1 inhibitor, was used. To inhibit fatty acid uptake via CD36 inhibition, we used Sulfosuccinimidyl Oleate (sodium salt), or SSO (Cayman Chemical # 1212012-37-7) at a concentration of 20 μM. For all drug treatments, MDMi media was changed on day 10 (after lentiviral knockdown) to remove virus from the media. After 3 days, inhibitors were added for 12-16 hours, and the next day (day 14) cells were assayed. DMSO was used as the vehicle control.

### Lipidomic Analysis

Samples were sent to the Lipidomics Core facility at Columbia University Medical Center. Lipids were extracted from equal amounts of material (1 million cells per sample). Lipid extracts were prepared via chloroform–methanol extraction, spiked with appropriate internal standards, and analyzed using a 6490 Triple Quadrupole LC/MS system (Agilent Technologies, Santa Clara, CA) as described previously [130].

### Statistical Analysis

All paired t-tests were 2-tailed, assuming normal (Gaussian) distribution. Multiple paired t-tests were performed for Fluidigm data analysis of knockdown MDMi, and an adjusted p-value of <0.05 (correcting for multiple comparisons using the Holm-Sidak method) was considered significant. Unadjusted p-values from multiple unpaired t-tests (2-tailed, Gaussian distribution) were used to analyze Fluidigm data for ALS patient-derived MDMi due to low sample numbers. DGAT-treated samples were compared to untreated samples using paired analyses. Statistical analysis on lipidomics data was performed using 2-tailed paired t-tests. All statistical analysis was performed in Prism.

## Acknowledgements

The authors are grateful to the participants of the New York Blood Center (NYBC) and the Eleanor and Lou Gehrig ALS Center at Columbia University for their contribution to this research. All participants were recruited and consented to participate and donate samples for research related to neuromuscular conditions under Columbia IRB Protocol # AAAK2000. This work was supported by the US National Institutes of Health grants R21AG073882, RF1AG058852, and R01AG076018 (EMB) and the Department of Defense grant AL200097 (WE). The content is solely the responsibility of the authors and does not necessarily represent the official views of the National Institutes of Health. EMB and WE are current Ludwig Scholars in the Carol and Gene Ludwig Center for Research in Neurodegeneration.

## Declaration of Competing Interest

The authors have no competing interests to declare.

## Author Contributions

K.K. and E.M.B. implemented the study and wrote the manuscript. E.A.G analyzed and interpreted lipidomics data. D.D. made PCA plots and heatmaps for Fluidigm data analysis. R.T helped with the analysis of confocal imaging data using CellProfiler. M.Y. and R.T. processed the NYBC and ALS blood samples used in the study. B.H., O.R. N.S. and W.E. supplied cryopreserved PBMCs from a thoroughly characterized ALS cohort. All authors read and edited the manuscript.

## Abbreviations List

ACAT1: Acyl-coenzyme A:cholesterol acyltransferase-1
AD: Alzheimer’s Disease
ALS: Amyotrophic lateral sclerosis
DAM: Disease-Associated Microglia
DGAT: Diacylglycerol Acyltransferase
LD: Lipid Droplet
MDMi: Monocyte-Derived Microglia-Like Cells

## Supplementary Figures

**Fig S1:** Western blot showing reduced protein levels in *TARDBP* knockdown. **A**) Western Blot images for scramble (Scr) vs TDP-43 knockdown (KD) showing TDP-43 band at ∼45KDa. Bottom panel shows GAPDH at ∼35KDa. Due to smaller TDP-43 bands at a similar size as GAPDH, minimal exposure was used to probe for GAPDH (used for normalization) **B**) Quantification of TDP-43 intensity normalized to GAPDH (N=6) **C**) TDP-43 nuclear to cytoplasmic ratio calculated using confocal imaging data (N=10). Statistical Analysis: paired t-test

**Fig S2:** Phospho-TDP-43 staining is unchanged in *TARDBP* knockdown. **A-C**) Mean intensity of nuclear, cytoplasmic and total phospho-TDP stain measured by CellProfiler quantification of confocal images (60X) for scramble compared to TDP-43 knockdown MDMi (N=8). **D)** Nuclear to cytoplasmic ratio of TDP-43 mean intensity (N=8) **E**) Confocal images of pTDP staining showing DAPI and Iba1 (60X). Scale bar = 10μm. Statistical Analysis: Paired t-test

**Fig S3:** IL1B expression and protein levels are reduced in knockdown MDMi. **A**)IL1B gene expression measured by qPCR in scramble vs TDP-43 knockdown MDMi (N=24). **B)** IL1B protein quantification using ELISA in scramble vs TDP-43 Knockdown MDMi (N=10) C) TREM2 protein quantification using ELISA in scramble vs TDP-43 KD (N=6). Statistical Analysis: Paired t-test

**Fig S4:** NBD-cholesterol uptake is not significantly upregulated in *TARDBP* knockdown stimulated with LPS compared to control. NBD-cholesterol mean intensity with LPS treatment: **A**) scramble compared to scramble+LPS (N=30). **B**) TDP-43 KD compared to TDP-43 KD+LPS (N=25) **C**) Ratio of Scramble+ LPS:Scr compared to TDP KD+LPS:TDP KD (N=25). **D**) Free cholesterol: Total cholesterol ratio in scramble versus TDP KD (N=8) **E**) Ratio of SUPs:Cells of total cholesterol in scramble versus TDP-43 KD (N=8). **F**) Mean Lipidtox intensity in scramble vs TDP-43 KD MDMi treated with ACAT inhibitor (N=5). Statistical Analysis: A-E) Paired t-test and F) 2-way ANOVA

**Fig S5:** Cell to supernatant ratio of triglycerides is increased in *TARDBP* knockdown MDMi. **A-C**) Cell to supernatant ratio of Triglyceride, total glycerol, and free glycerol in scramble versus *TARDBP* knockdown (TDP-43 KD) MDMi (N=11). Statistical Analysis: Paired t-test

**Fig S6:** ACAT1 inhibitor does not alter triglyceride levels in MDMi. **A-C**): Total glycerol, Free Glycerol, and Triacylglycerol levels in cell lysates of scramble versus *TARDBP* knockdown (TDP-43 KD) treated with and without ACAT1 inhibitor, measured using Promega TriGlo Assay. **D**) Total glycerol measured in supernatants of scramble versus TDP-43 knockdown MDMi treated with and without ACAT1 inhibitor (Free glycerol levels were too low to quantify) (N=12). Statistical analysis: 2-Way ANOVA

**Fig S7:** qPCR validation of expression of triglyceride-associated genes. **A-D**) Gene expression measured using TaqMan qPCR in scramble versus *TARDBP* knockdown (TDP-43 KD) MDMi (N=15). Statistical Analysis: Paired t-test

**Fig S8:** DGAT inhibitor significantly alters gene expression in control and *TARDBP* knockdown MDMi. Heatmaps showing Fluidigm data for gene expression alterations in control vs *TARDBP* knockdown (KD), along with effect of DGAT inhibitor on control and *TARDBP* knockdown. Fold change is expressed as Log2FC. Genes that were significantly altered (p-value<0.05) are indicated by the black dot and grey dot indicates genes that were close to significance. Statistical analysis: Multiple paired t-test.

**Fig S9:** Fatty acid uptake is altered by CD36 inhibition, but lipid droplets are not. **A**) C12-BODIPY mean intensity measured by CellProfiler using confocal images for scramble and TDP-43 KD MDMi treated with DGAT inhibitors overnight (N=4). **B**) C12-BODIPY mean intensity measured by CellProfiler using confocal images, for scramble and TDP-43 KD MDMi treated with SSO for 4 hours (N=7). **C**) LipidTox mean intensity measured by CellProfiler using confocal images for scramble and TDP-43 KD cells +/- SSO for 24 hours (N=5). **D**) LipidTox mean intensity for scramble vs KD treated with SSO for 24 hours (N=8). Statistical Analysis: 2 Way ANOVA

**Fig S10:** Lipidomic analysis. **A-E**) Heat maps showing lipidomic analysis of certain species: A) Acylcarnitines B) Triglycerides C) Monohexylceramides D) Bis(monoacylglycero)phosphate, and E) Lactosylceramides. Bold species were significant (paired t-test, p<0.05, N=4)

**Fig S11:** ALS patient-derived MDMi demographics and mean lipid droplet quantification. **A**) Demographics of TDP-ALS mutant patients and age and sex matched control samples used to obtain PBMCs for MDMi. **B**) TARDBP gene expression measured by TaqMan qPCR in control vs TDP-ALS mutant patients (N=3). **C**) Confocal images showing LipidTox staining in controls and TDP-ALS mutant MDMi with DGAT inhibitor treatment. **D**) Quantification of mean LipidTox intensity using imaging data from CellProfiler in control and TDP-ALS MDMi +/- DGAT inhibitors. **E**) Quantification of mean lipid droplet (LD) area by CellProfiler in scramble and TDP-ALS MDMi +/- DGAT inhibitors. **F**) Quantification of mean lipid droplet (LD) radius using CellProfiler in scramble and TDP-ALS MDMi +/- DGAT inhibitors. **G**) Confocal Images showing LipidTox staining in control vs sALS patient MDMi. **H**) LipidTox mean intensity in control and sALS MDMi quantified using CellProfiler (N=4). **I**) Mean lipid droplet (LD) area in control and sALS MDMi quantified using CellProfiler (N=3). Statistical analysis D-F) 2-Way ANOVA, B, H, I) Unpaired t-test.

**Fig S12:** TDP-ALS samples cluster together whereas sporadic ALS and controls are heterogenous. **A)** Clustered heat map using z-score matrix and row clustering for controls, TDP-ALS and sALS samples. **B**) PCA plot for ALS-TDP and matched controls (N=3), and **C**) PCA plot for sALS and matched controls (N=6). **D-E**) Volcano plots showing Log2FC versus -Log10P for Fluidigm gene expression data in ALS-TDP MDMi compared to controls and sALS MDMi compared to controls. Log2FC calculated using average expression values for each group. Statistical test: unpaired multiple t-test (unadjusted p-values).

**Table 1:**
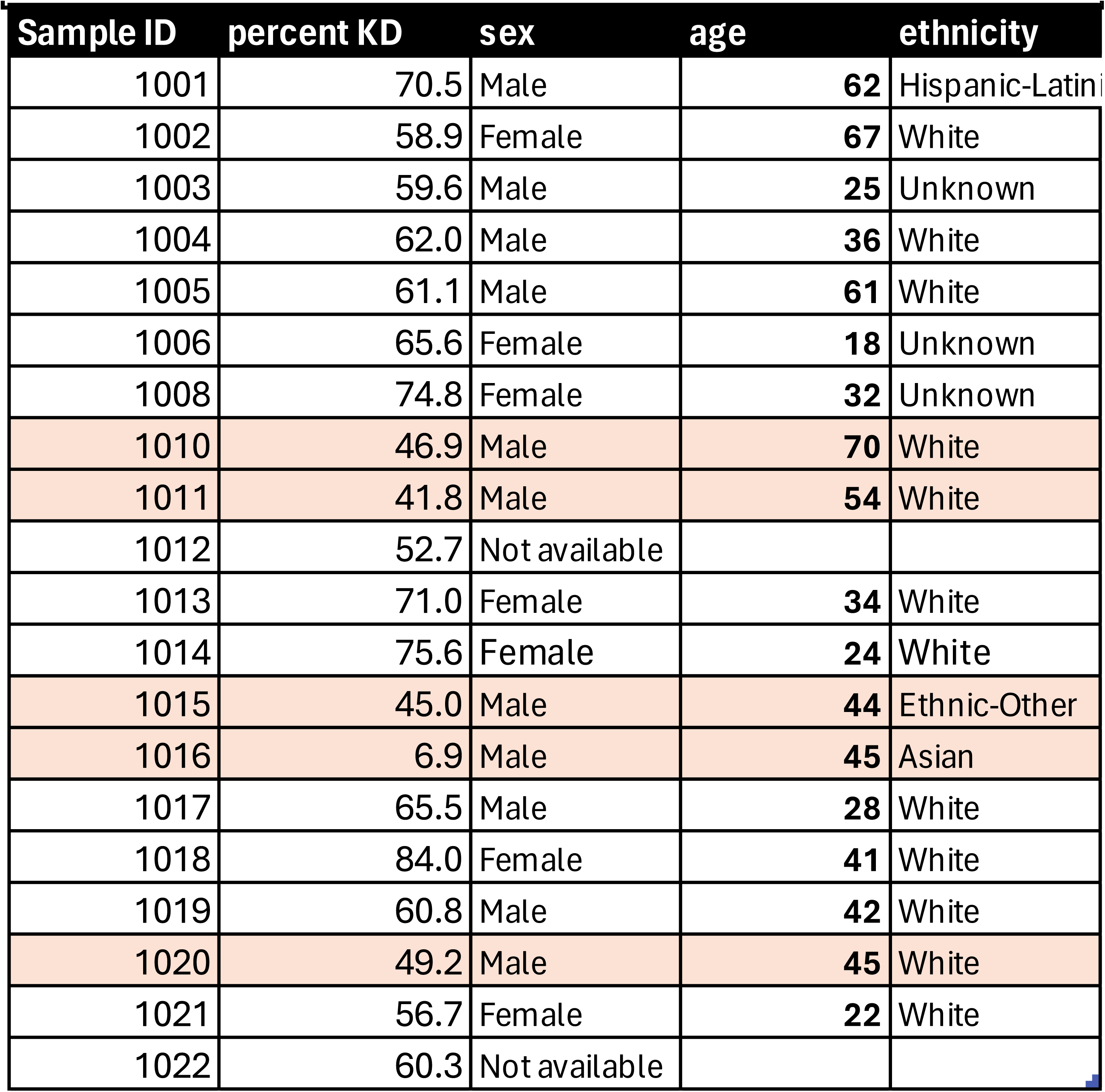
Demographics of NYBC samples used for Fluidigm nalysis (highlighted samples were removed from final data due o low KD efficiency)

| Sample ID | percent KD | sex | age | ethnicity |
| --- | --- | --- | --- | --- |
| 1001 | 70.5 | Male | 62 | Hispanic-Latini |
| 1002 | 58.9 | Female | 67 | White |
| 1003 | 59.6 | Male | 25 | Unknown |
| 1004 | 62.0 | Male | 36 | White |
| 1005 | 61.1 | Male | 61 | White |
| 1006 | 65.6 | Female | 18 | Unknown |
| 1008 | 74.8 | Female | 32 | Unknown |
| 1010 | 46.9 | Male | 70 | White |
| 1011 | 41.8 | Male | 54 | White |
| 1012 | 52.7 | Not available |  |  |
| 1013 | 71.0 | Female | 34 | White |
| 1014 | 75.6 | Female | 24 | White |
| 1015 | 45.0 | Male | 44 | Ethnic-Other |
| 1016 | 6.9 | Male | 45 | Asian |
| 1017 | 65.5 | Male | 28 | White |
| 1018 | 84.0 | Female | 41 | White |
| 1019 | 60.8 | Male | 42 | White |
| 1020 | 49.2 | Male | 45 | White |
| 1021 | 56.7 | Female | 22 | White |
| 1022 | 60.3 | Not available |  |  |

**Table 2:** Demographics of sporadic ALS patients and NYBC control samples.

|  |  |  |  |  |  |
| --- | --- | --- | --- | --- | --- |
| Sporadic ALS Patients |  |  | Matched NYBC Control |  |  |
| ID | Sex | Age at Cx | ID | Sex | Age |
| sALS-1 | Male | 39 | 1028 | Male | 54 |
| sALS-2 | Male | 54 | 1005 | Male | 61 |
| sALS-3 | Female | 53 | 1026 | Female | 63 |
| sALS-4 | Male | 48 | 1020 | Male | 45 |
| sALS-5 | Male | 64 | 1005 | Male | 61 |
| sALS-6 | Female | 38 | 1024 | Female | 40 |
| sALS-7 | Female | 59 | 1023 | Female | 52 |
| sALS-8 | Male | 58 | 1027 | Male | 46 |
| sALS-9 | Female | 55 | 1025 | Male | 40 |

## Notes

### Competing Interest Statement

The authors have declared no competing interest.

