## Supplementary figures and images for "Loss of Nuclear TDP-43 Impairs Lipid Metabolism in Microglia-Like Cells"

### Supplemental Figure 1

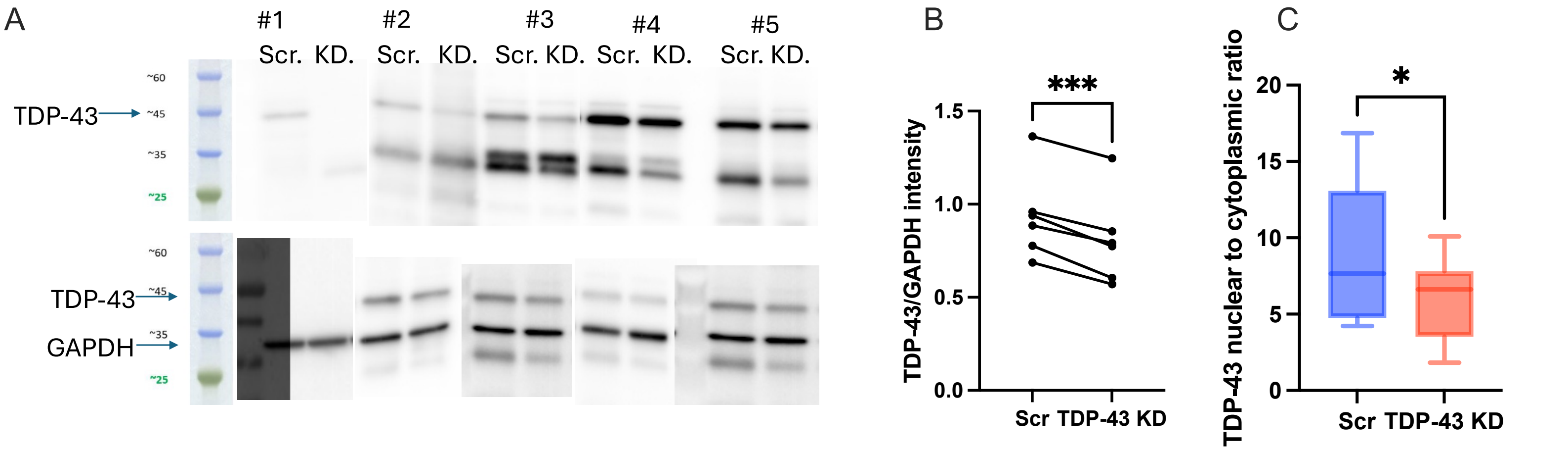

### Supplemental Figure 2

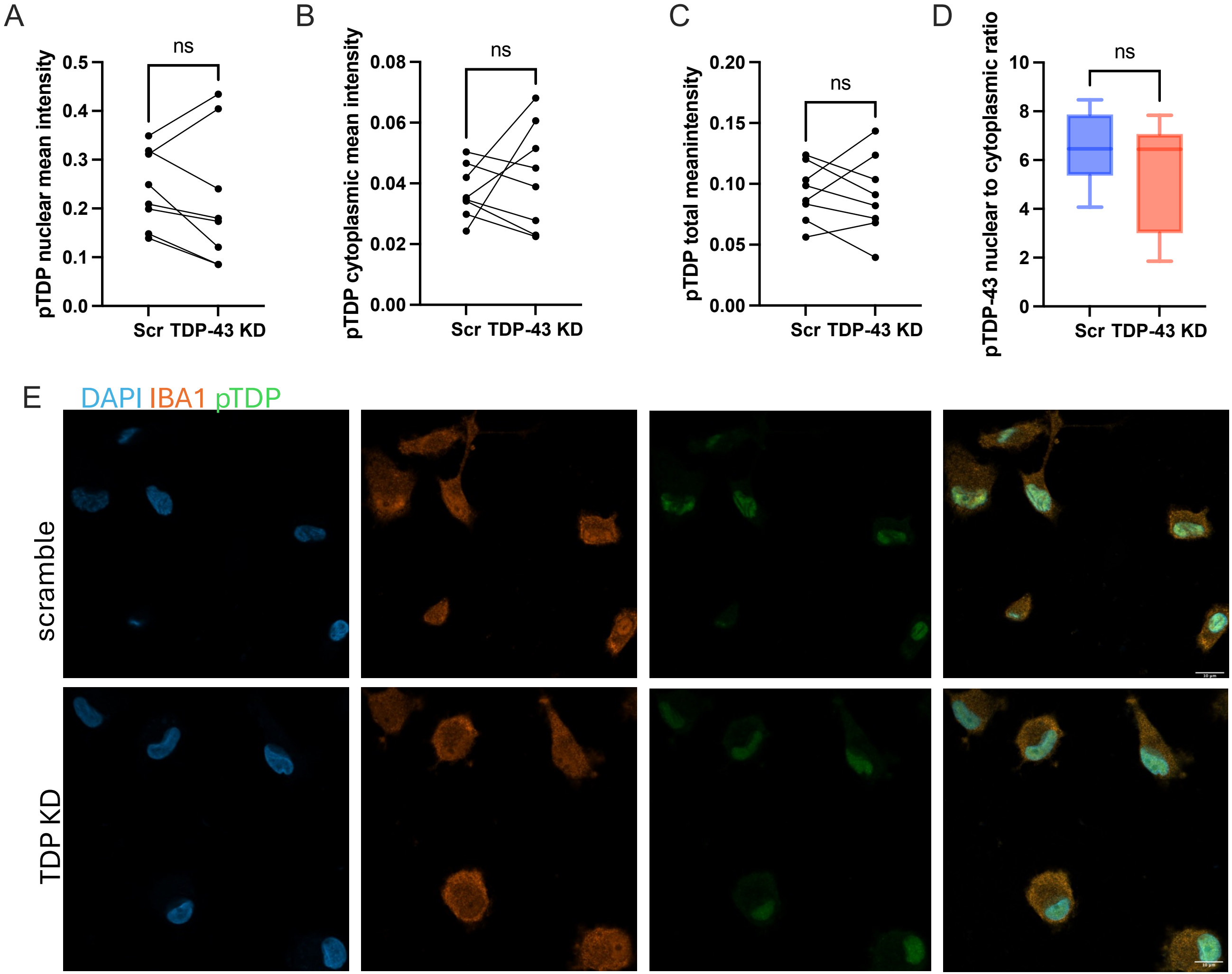

### Supplemental Figure 3

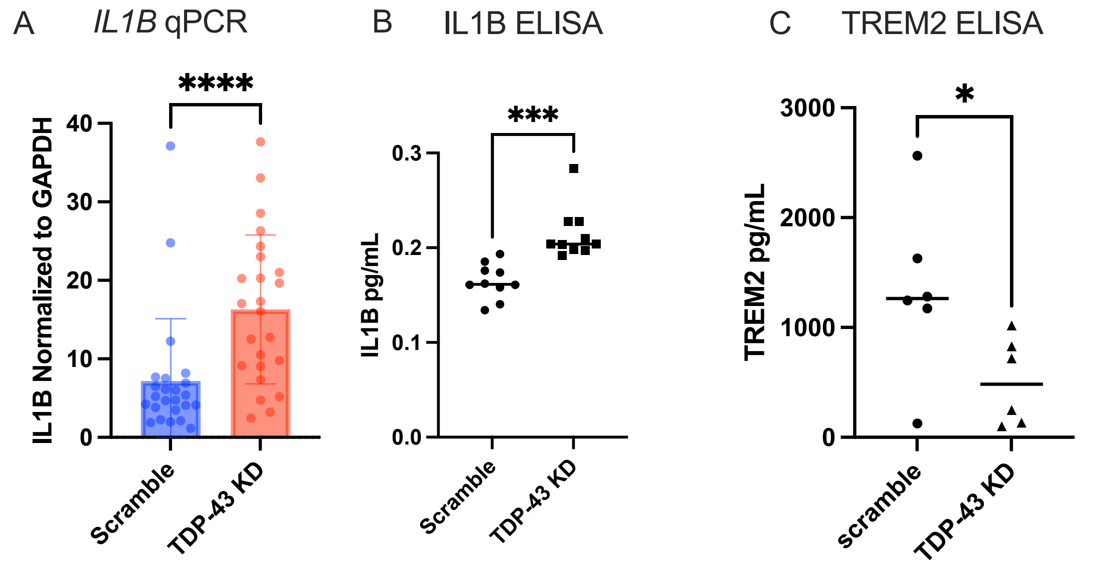

### Supplemental Figure 4

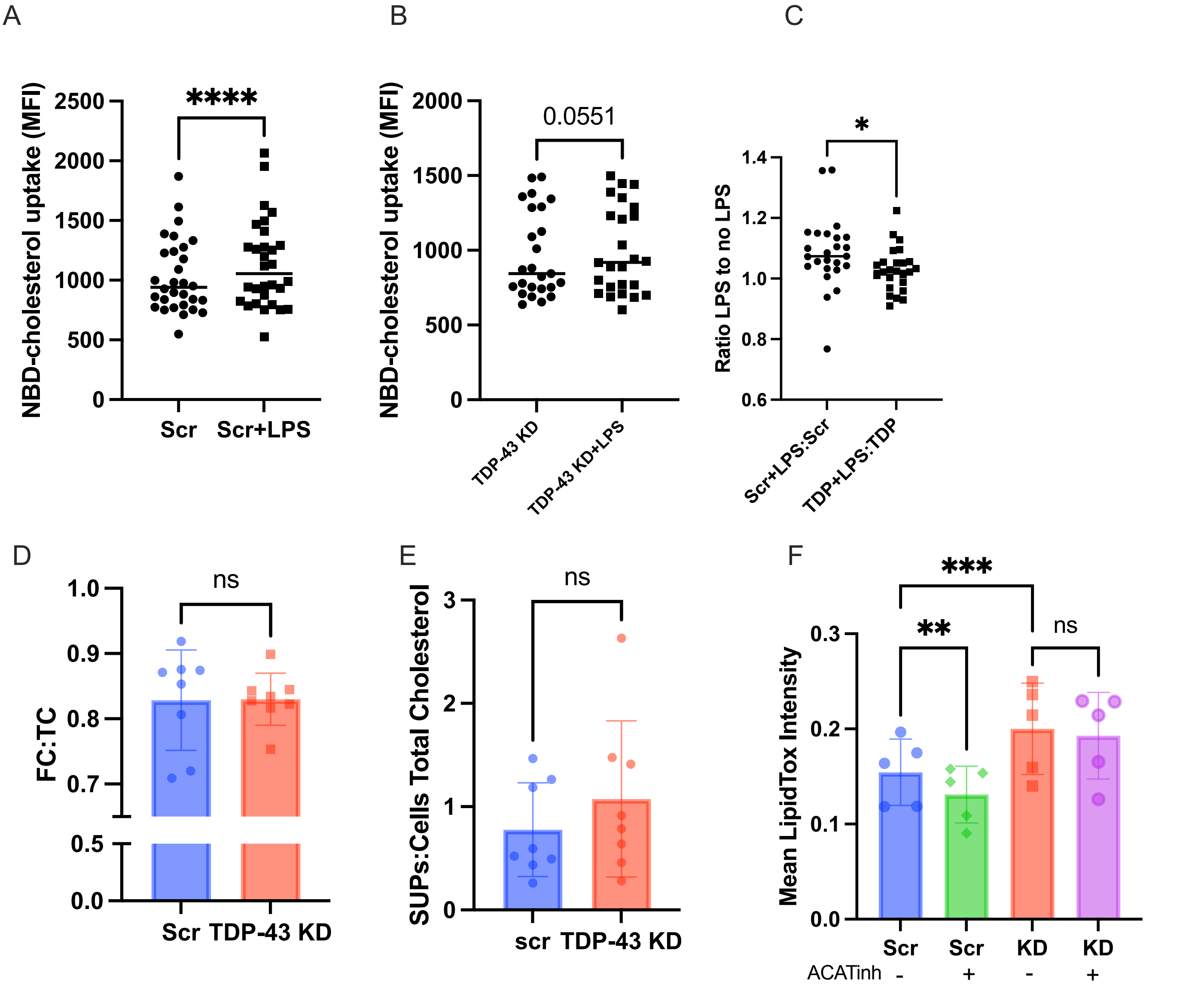

### Supplemental Figure 5

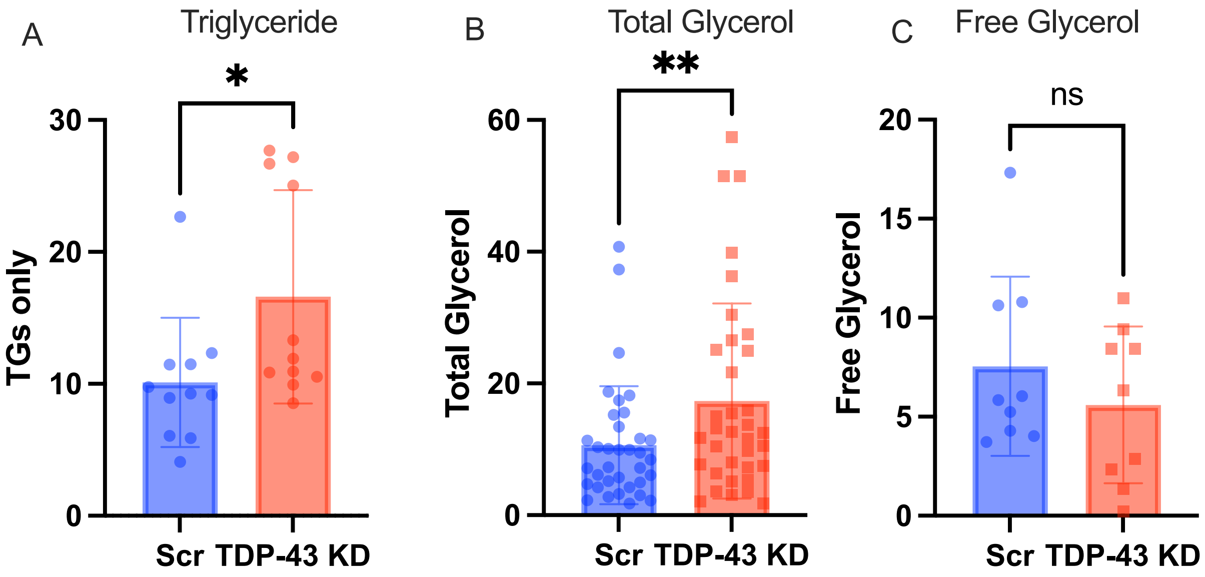

### Supplemental Figure 7

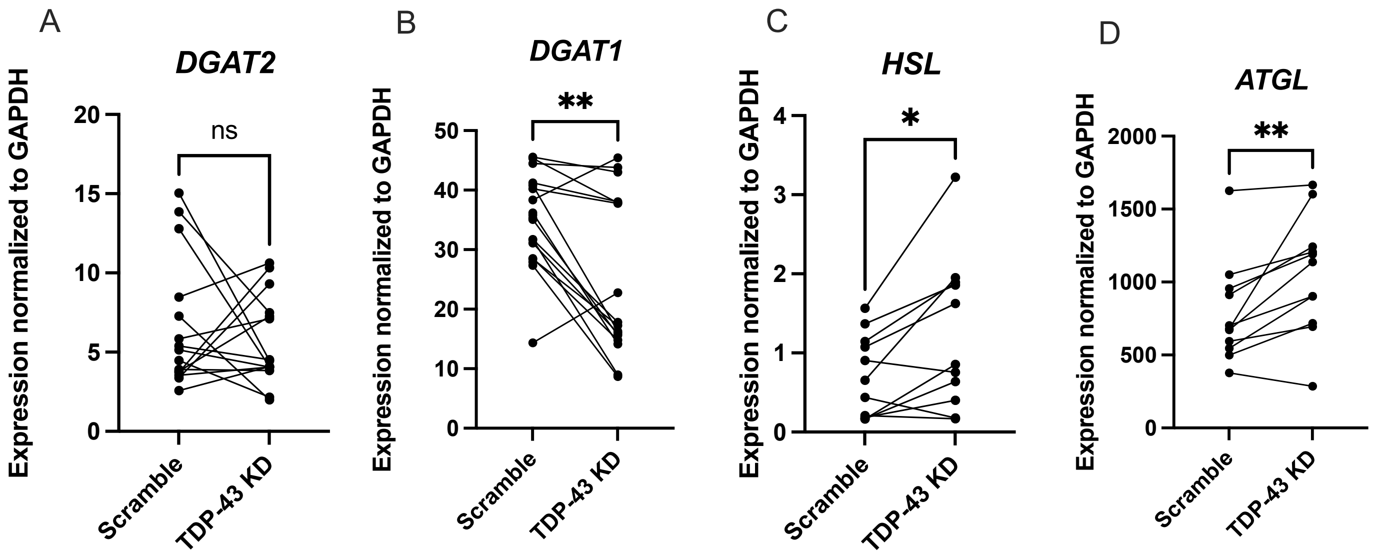

### Supplemental Figure 9

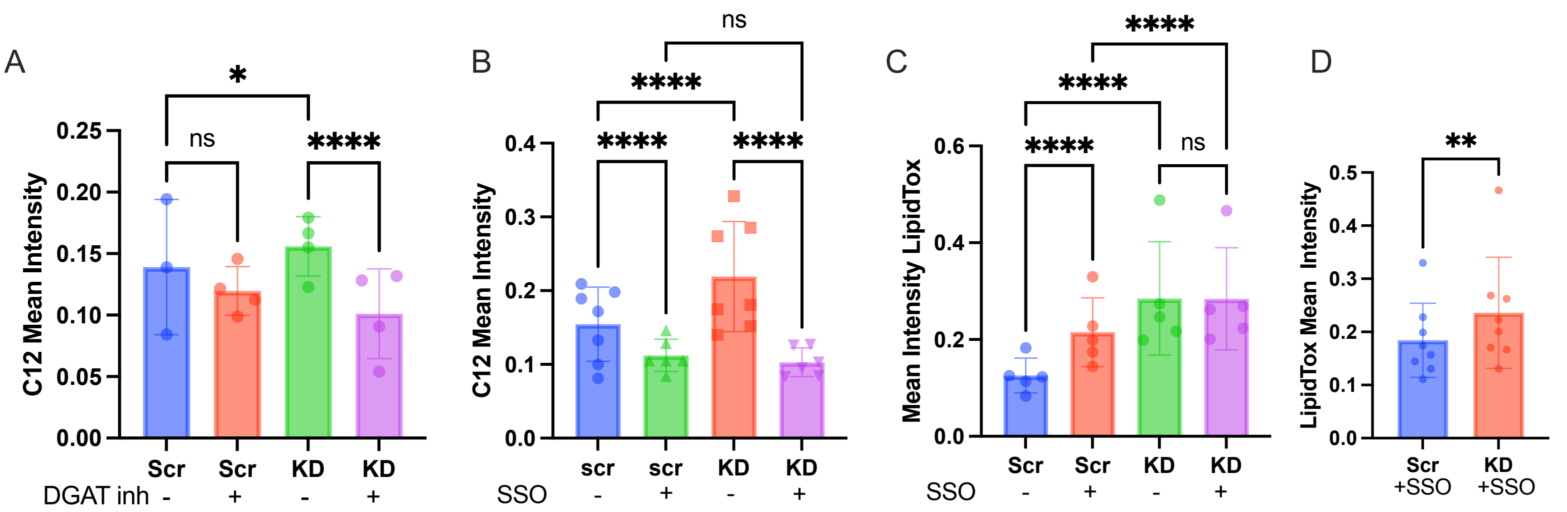

### Supplemental Figure 11

A

| ID | Age | Sex | Variant |
|---|---|---|---|
| TARDBP-1 | 47 | M | I383V |
| TARDBP-2 | 43 | M | A382T |
| TARDBP-3 | 41 | F | S379P |
| Control-1 | 45 | F | NA |
| Control-2 | 46 | M | NA |
| Control-3 | 45 | M | NA |

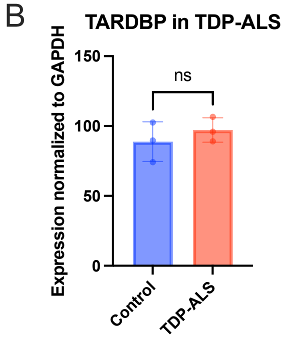


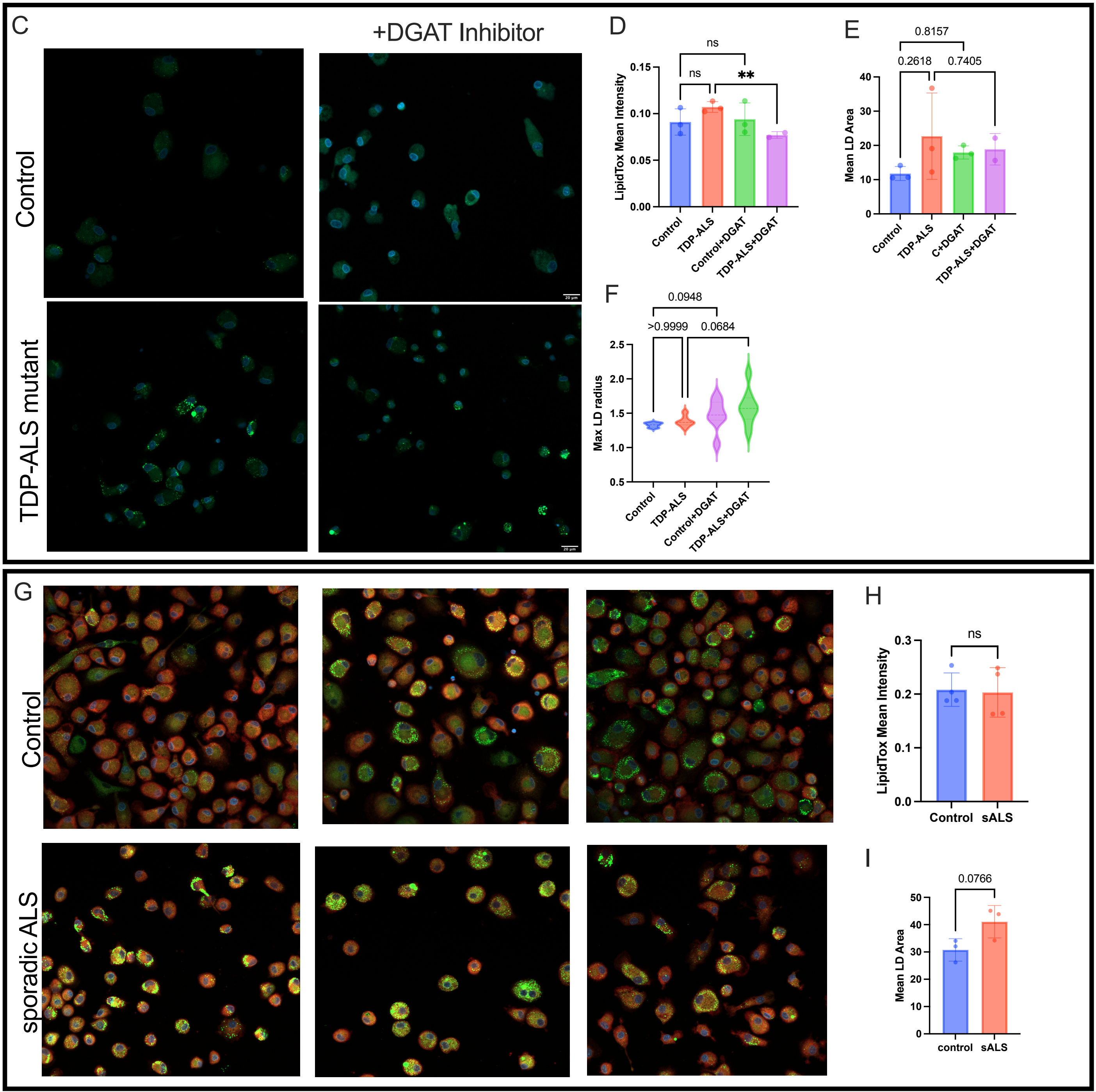

### Supplemental Figure 12

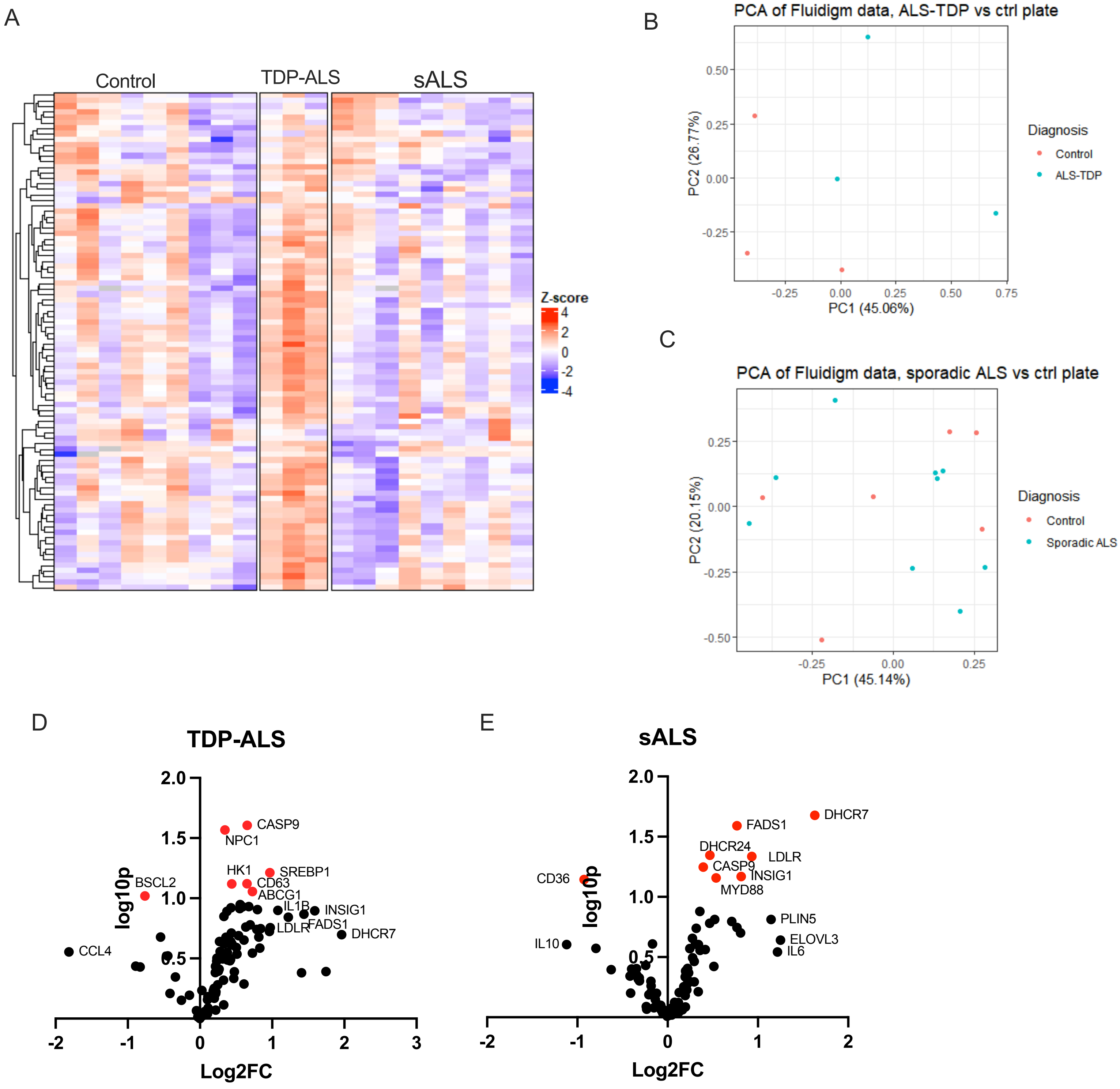
