## Supplemental Figure 10 for "Loss of Nuclear TDP-43 Impairs Lipid Metabolism in Microglia-Like Cells"

| Species | Log2FC | p-value |
|---|---|---|
| **AC C12:0** | **-0.4926895** | **0.01473** |
| AC C18:0 | -0.3819312 | 0.124576 |
| AC C18:1 | -0.0272371 | 0.959943 |
| AC C14:0 | 0.06807314 | 0.535481 |
| AC C16:0 | 0.09911742 | 0.565382 |
| AC C8:0 | 0.09974797 | 0.117706 |
| AC C2:0 | 0.24505412 | 0.235139 |
| AC C6:0 | 0.27510206 | 0.185596 |
| AC C3:0 | 0.41972727 | 0.261998 |

| Species | Log2FC | p-value |
|---|---|---|
| TG 52:0/18:0 | -0.4647029 | 0.088086 |
| **TG 54:0/18:0** | **-0.4379092** | **0.041199** |
| TG 54:1/18:0 | -0.4187514 | 0.090482 |
| TG 52:1/18:0 | -0.2561546 | 0.260001 |
| TG 50:0/16:0 | -0.2297343 | 0.298548 |
| TG 60:8/22:6 | -0.1503512 | 0.353387 |
| TG 56:4/20:4 | -0.1212539 | 0.391002 |
| TG 54:2/18:0 | -0.1071201 | 0.472167 |
| TG 56:6/20:4 | 0.10114567 | 0.702278 |
| TG 48:1/16:0 | 0.11161402 | 0.456214 |
| TG 56:5/20:4 | 0.11560761 | 0.773277 |
| TG 54:5/20:4 | 0.13691029 | 0.466328 |
| TG 54:4/20:4 | 0.13919438 | 0.617294 |
| TG 58:6/20:4 | 0.14450665 | 0.538518 |
| TG 60:9/22:6 | 0.15155263 | 0.091721 |
| TG 60:7/22:6 | 0.16088575 | 0.637618 |
| TG 54:4/18:1 | 0.16392712 | 0.197116 |
| TG 58:7/20:4 | 0.18369464 | 0.604959 |
| TG 56:5/18:1 | 0.18489887 | 0.375954 |
| TG 50:3/16:1 | 0.18707436 | 0.109446 |
| TG 54:7/18:1 | 0.1872822 | 0.590801 |
| TG 56:7/20:4 | 0.20219151 | 0.56317 |
| TG 52:4/18:1 | 0.20239439 | 0.197012 |
| TG 54:5/18:1 | 0.23499935 | 0.173283 |
| TG 54:6/18:1 | 0.25741814 | 0.326142 |
| TG 52:5/20:4 | 0.26684938 | 0.297641 |
| TG 52:5/18:1 | 0.29151204 | 0.23832 |
| TG 54:6/20:4 | 0.30291245 | 0.279336 |
| TG 56:8/20:4 | 0.383049 | 0.489397 |
| TG 54:7/20:4 | 0.46191965 | 0.180362 |
| TG 56:9/20:4 | 0.57029964 | 0.235139 |

| Species | Log2FC | p-value |
|---|---|---|
| BMP 30:0 | -0.4519877 | 0.18169 |
| BMP 38:0 | -0.4131485 | 0.391002 |
| BMP 34:0 | -0.4125906 | 0.141122 |
| BMP 32:0 | -0.3622026 | 0.391002 |
| BMP 40:6 | -0.3340155 | 0.339736 |
| BMP 40:7 | -0.3264576 | 0.23054 |
| BMP 38:6 | -0.3036856 | 0.435474 |
| BMP 34:1 | -0.2932943 | 0.180759 |
| BMP 40:5 | -0.2384883 | 0.391002 |
| BMP 36:1 | -0.236797 | 0.164419 |
| BMP 38:1 | -0.2158493 | 0.091721 |
| BMP 36:0 | -0.2097623 | 0.252215 |
| BMP 36:2 | -0.2020164 | 0.055853 |
| BMP 42:5 | 0.11019075 | 0.391002 |
| BMP 42:6 | 0.14519742 | |
| BMP 42:7 | 0.37303084 | 0.391002 |

| Species | Log2FC | p-value |
|---|---|---|
| MhCer d18:0/18:0 | -0.3830938 | 0.296689 |
| MhCer d18:0/20:0 | -0.274147 | 0.495025 |
| MhCer d18:0/22:0 | -0.1473258 | 0.256007 |
| MhCer d18:0/26:1 | 0.21843098 | 0.391002 |
| MhCer d18:0/18:1 | 0.2733287 | 0.465148 |
| **MhCer d18:1/26:0** | **0.27548198** | **0.040519** |
| MhCer d18:1/24:1 | 0.41162286 | 0.106367 |
| MhCer d18:1/26:1 | 0.44084378 | 0.013216 |
| MhCer d18:1/20:1 | 0.46908398 | 0.264649 |
| MhCer d18:1/22:1 | 0.51844721 | 0.183213 |
| MhCer d18:1/18:1 | 0.57408025 | 0.354283 |
| MhCer d18:0/16:1 | 0.58274295 | 0.304289 |
| MhCer d18:1/16:0 | 0.58615095 | 0.292021 |
| MhCer d18:1/16:1 | 0.6182083 | 0.291888 |

| Species | Log2FC | p-value |
|---|---|---|
| LacCer d18:0/18:0 | -1.2642494 | 0.294128 |
| LacCer d18:0/20:1 | -1.2458338 | 0.223427 |
| LacCer d18:0/20:0 | -1.15507 | 0.229983 |
| LacCer d18:0/22:0 | -1.0981867 | 0.244842 |
| LacCer d18:0/22:1 | -1.0870795 | 0.242454 |
| LacCer d18:0/18:1 | -1.026333 | 0.223427 |
| LacCer d18:0/24:1 | -1.0152604 | 0.241732 |
| LacCer d18:0/16:0 | -0.9872804 | 0.244645 |
| LacCer d18:0/24:0 | -0.954114 | 0.197849 |
| LacCer d18:1/18:0 | -0.8832978 | 0.213901 |
| LacCer d18:0/26:1 | -0.7370053 | 0.277873 |
| LacCer d18:1/20:0 | -0.671897 | 0.221175 |
| LacCer d18:0/16:1 | -0.6509261 | 0.157218 |
| LacCer d18:1/16:0 | -0.5284937 | 0.118322 |
| LacCer d18:0/26:0 | -0.5063223 | 0.21517 |
| LacCer d18:1/24:0 | -0.500989 | 0.072238 |
| LacCer d18:1/24:1 | -0.4867291 | 0.142028 |
| LacCer d18:1/20:1 | -0.4715089 | 0.09218 |
| LacCer d18:1/26:0 | -0.460357 | 0.073505 |
| LacCer d18:1/26:1 | -0.4580819 | 0.203053 |
| LacCer d18:1/18:1 | -0.4326015 | 0.061684 |
| LacCer d18:1/22:0 | -0.419072 | 0.146256 |
| LacCer d18:1/22:1 | -0.4123798 | 0.131241 |
| LacCer d18:1/16:1 | -0.3821275 | 0.072942 |
