## Supplemental Figure 8 for "Loss of Nuclear TDP-43 Impairs Lipid Metabolism in Microglia-Like Cells"

**Legend:** Log₂ Fold Change: -3, -2, -1, 0, 1, 2, 3 p-value: ● p < 0.01 • p < 0.05

| Category | Gene | C vs KD | KD vs KD+DGATinh | C vs C+DGAT inh | C+DGATinh vs KD+DGATinh |
|---|---|---|---|---|---|
| Cholesterol Metabolism (23) | INSIG1 | **-2.0** ● | -0.9 | -0.8 | **-2.1** ● |
| | ABCC4 | **-1.2** ● | **-1.1** ● | -2.0 | -0.6 · |
| | MBTPS1 | **-1.2** ● | -0.6 · | -0.1 | **-1.7** ● |
| | SREBP2 | **-0.9** ● | **-1.3** ● | -1.2 | **-1.0** ● |
| | LSS | **-0.9** ● | **-0.9** • | -0.9 | **-0.9** ● |
| | SCARB1 | **-0.8** ● | **-1.2** • | -1.3 | **-0.9** ● |
| | ABCA1 | **-0.7** ● | -1.7 | -1.1 | **-1.4** ● |
| | LCAT | **-0.7** • | -0.2 | -0.1 | -0.8 · |
| | LDLR | -0.6 | -0.7 · | -0.3 | **-1.2** ● |
| | NPC1 | **-0.5** ● | **-1.5** ● | -0.8 | **-1.2** ● |
| | DHCR24 | **-0.5** • | **-2.0** ● | -0.9 | **-1.3** ● |
| | MBTPS2 | **-0.5** • | **-1.1** ● | -0.5 | **-1.1** ● |
| | HMGCR | **-0.5** • | **-1.1** • | -1.0 | **-0.8** • |
| | CYP27A1 | **-0.3** • | -1.2 | -1.6 | 0.1 |
| | LRP1 | **-0.3** ● | **-1.5** • | -1.8 | 0.1 · |
| | SC5D | -0.1 | -1.0 | -1.6 · | 0.2 |
| | ACAT1 | -0.0 | **-1.6** ● | -0.9 | **-0.8** ● |
| | ERLIN2 | -0.0 | -1.0 | -0.8 | -0.1 |
| | LXR | 0.0 | -0.1 | -0.4 | 0.4 |
| | ABCG1 | 0.0 | -0.5 | -0.7 | 0.2 |
| | DHCR7 | 0.3 | **-1.7** ● | -0.8 · | -0.2 |
| | SOAT1 | **0.4** ● | **-1.4** • | -1.1 | 0.1 |
| | STARD4 | **0.6** ● | -0.4 | -0.2 | 0.5 |
| Fatty Acid Metabolism (15) | ELOVL7 | **-1.5** ● | 0.5 · | 0.5 | -0.9 · |
| | PLA2G7 | **-0.7** ● | **-1.1** ● | -0.9 | **-0.8** ● |
| | Asah1 | **-0.6** ● | -1.2 · | -1.1 | **-0.7** • |
| | ACSL5 | -0.5 · | -0.3 | -0.4 | -0.6 |
| | CPT2 | **-0.4** ● | **-1.7** • | -1.1 | **-0.7** ● |
| | FASN | -0.3 | **-1.5** ● | **-0.9** ● | **-0.9** • |
| | CD36 | **-0.3** • | -1.1 · | -0.9 | **-0.6** • |
| | SREBP1 | -0.0 | **-0.3** ● | -0.1 | **-0.3** • |
| | PPARG | 0.1 | -0.7 | -0.6 | -0.0 |
| | FABP5 | 0.2 · | 0.2 | 0.2 | 0.3 |
| | ACSL1 | **0.5** ● | -0.6 | -0.5 | 0.4 · |
| | ACSL4 | **0.5** ● | **-1.4** • | -0.7 | -0.2 |
| | FADS1 | **0.6** ● | **-1.4** ● | -1.0 | 0.3 |
| | FABP4 | **3.0** ● | 0.5 | 1.1 | **2.3** • |
| | ELOVL3 | **3.5** • | 0.1 | **1.8** • | 1.3 |
| Triglyceride Metabolism (18) | LPL | **-1.7** ● | **-2.3** ● | -1.4 | **-2.5** ● |
| | GPAT3 | **-1.4** ● | **-1.6** ● | -1.3 | **-1.6** ● |
| | LIPA | **-1.3** ● | -1.6 · | -1.3 | **-1.5** ● |
| | MGAT1 | **-1.1** ● | 0.1 | 0.0 | **-1.0** ● |
| | LPCAT1 | **-1.0** ● | -0.9 · | -0.5 | **-1.3** • |
| | APOBR | **-0.7** ● | **-2.4** ● | -1.8 · | **-1.0** ● |
| | AGPAT1 | **-0.7** ● | **-1.5** ● | -1.0 | **-0.9** ● |
| | DDHD2 | **-0.6** ● | -1.0 · | -0.8 | **-0.8** • |
| | PLIN3 | **-0.5** ● | **-1.8** ● | -1.3 | **-1.0** ● |
| | DGAT1 | **-0.3** ● | -0.5 | -0.2 | **-0.5** • |
| | AGPAT2 | -0.2 | 0.1 | 0.3 | -0.7 |
| | MGLL | -0.1 | -0.5 | -0.7 | 0.2 |
| | DGAT2 | -0.1 | **-0.7** • | -0.7 | -0.4 |
| | GPAT4 | -0.1 | **-1.2** • | -1.0 | -0.5 |
| | PLIN2 | **0.3** • | **0.5** • | **0.3** • | 0.6 |
| | ATGL | **0.5** ● | 0.4 · | **0.2** • | **0.6** • |
| | HSL (LIPE) | 1.4 · | -1.4 | 1.3 | 0.1 |
| | BSCL2 | **4.2** ● | **-1.3** • | 0.7 | **3.9** ● |

| Category | Gene | C vs KD | KD vs KD+DGATinh | C vs C+DGAT inh | C+DGATinh vs KD+DGATinh |
|---|---|---|---|---|---|
| Carbohydrate Metabolism (14) | PFKP | **-1.7** ● | -0.2 | -0.5 | **-1.3** ● |
| | G6PD | **-1.3** ● | -0.5 | -0.3 | **-1.5** ● |
| | GLUT1 | **-1.1** ● | -0.4 | -0.3 · | **-1.1** • |
| | ENO1 | **-0.8** ● | **-1.0** ● | **-0.5** • | **-1.3** ● |
| | PC | **-0.6** ● | -0.4 | -0.8 | 0.0 |
| | TPI1 | **-0.5** • | **0.6** • | 0.2 | -0.1 |
| | PDH | **-0.5** ● | **-1.4** ● | -1.7 | -0.2 |
| | HK2 | -0.4 · | -0.1 | 0.2 | -0.7 |
| | PGK1 | **-0.4** ● | **-0.7** ● | **-0.6** ● | **-0.5** • |
| | GPI | **-0.3** ● | **-0.7** • | -0.4 | **-0.7** ● |
| | PKM | -0.3 · | **-0.9** ● | **-0.5** ● | **-0.7** • |
| | ALDOA | 0.1 | **-1.6** • | -1.2 | -0.4 |
| | ALDOC | 0.2 | **1.2** • | 0.4 | **0.7** • |
| | HK1 | **0.9** ● | **-0.9** • | -0.5 | **0.4** ● |
| Immune Response (25) | TREM2 | **-1.9** ● | **-2.1** ● | -1.4 | **-2.6** ● |
| | IL10 | **-1.7** ● | -0.9 | 0.1 | **-3.1** ● |
| | TLR4 | **-1.6** ● | -0.6 | -0.4 | **-1.5** ● |
| | CD9 | **-0.9** ● | -1.8 | -1.9 · | -0.8 |
| | CXCL1 | **-0.8** • | -2.1 | -0.9 | **-3.8** ● |
| | PLCG2 | **-0.8** ● | -0.7 | -0.9 | **-0.6** ● |
| | IRF7 | **-0.7** ● | 0.2 | -0.2 | -0.1 |
| | CD33 | **-0.6** ● | -1.3 · | -1.9 | **-0.1** • |
| | CD68 | **-0.4** ● | -1.2 | -1.0 | -0.6 · |
| | INPP5D | -0.3 | **-2.2** ● | -1.1 | **-1.2** ● |
| | ISG15 | -0.2 | **0.5** • | 0.3 · | -0.0 |
| | CD63 | -0.2 | **-1.9** ● | -0.7 | **-1.6** • |
| | TNFA | -0.2 | **-2.5** ● | -1.0 | -1.7 |
| | MYD88 | -0.2 · | **-0.9** ● | -0.6 | **-0.4** • |
| | IL18 | -0.1 | **-1.8** ● | -0.9 | **-0.8** ● |
| | IFNB1 | -0.0 | **0.7** • | **0.6** ● | -0.1 |
| | DAP12 | 0.0 | **-1.6** ● | -1.3 | -0.2 |
| | IFNA1 | 0.0 | **0.7** • | **0.8** ● | -0.0 |
| | IFIT3 | 0.1 | 0.2 | 0.4 | -0.1 |
| | CCL5 | 0.3 | -0.3 | 0.7 | -0.6 |
| | CXCL8 | 0.3 | 0.0 | 1.2 · | -0.8 |
| | NLRP3 | 0.3 | -0.8 | -0.5 | 0.6 |
| | IL1B | **1.6** ● | **-1.5** ● | 0.1 | -0.3 |
| | TREM1 | **1.7** ● | -1.0 | -0.6 | 0.8 |
| | CCL4 | **2.4** ● | **-3.7** ● | 0.3 | -1.0 |
| Phospholipid Metabolism (8) | PLA2G15 | **-0.9** ● | -0.8 | -1.4 | **-0.5** • |
| | PISD | **-0.6** ● | -0.4 | -0.1 | **-0.9** ● |
| | SPTLC1 | **-0.6** ● | -0.9 | -1.1 | -0.4 |
| | CERS1 | -0.5 | **1.1** • | 0.4 | 0.2 |
| | PLD1 | **-0.3** • | -1.2 | -0.9 | -0.6 |
| | STARD7 | -0.2 | **-1.1** ● | -0.5 | **-0.8** • |
| | EDG3 | -0.2 | **0.8** ● | 0.4 | **0.3** • |
| | SPTLC2 | -0.1 | **-1.3** • | **-2.3** • | 0.8 |
| Autophagy (3) | ATG16L1 | **-0.7** ● | **-1.3** • | -0.7 | **-1.5** ● |
| | CTSD | **-0.6** ● | -1.2 | -1.6 | -0.3 |
| | ATG7 | **-0.5** ● | **-1.1** • | **-1.1** • | -0.4 |
| Other (5) | TARDBP | **-1.5** ● | -0.7 · | -0.8 | **-1.4** ● |
| | FUS | **-1.0** ● | **-1.4** ● | -1.0 | **-1.3** ● |
| | CASP9 | **-0.8** ● | -0.2 | 0.2 | **-1.1** ● |
| | ATP5F1 | **-0.7** ● | **-1.5** ● | -1.4 · | **-0.7** • |
| | ALDH1A2 | **-0.6** • | 0.9 | 0.7 | 1.4 |
