## Supplemental Figure 6 for "Loss of Nuclear TDP-43 Impairs Lipid Metabolism in Microglia-Like Cells"

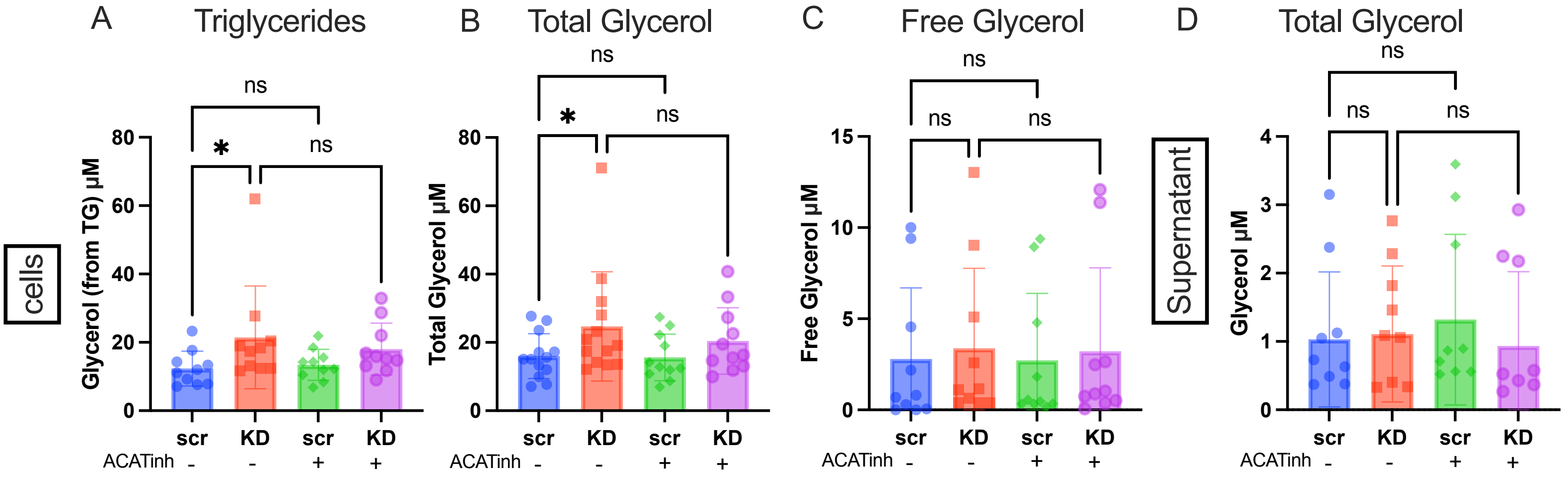


Fig S6A-C): Total glycerol, Free Glycerol and Triacylglycerol levels in cell lysates of scramble vs TDP-43 knockdown treated with and without ACAT1 inhibitor, measured using Promega TriGlo Assay. D) Total glycerol measured in supernatants of scramble vs TDP-43 knockdown MDMi treated with and without ACAT1 inhibitor (Free glycerol levels were too low to quantify)
Statistical analysis: 2-Way ANOVA
